# Identification of optimal fluorophores for use in the *Drosophila* embryo

**DOI:** 10.1101/2025.09.29.678560

**Authors:** Bernardo Chapa-y-Lazo, Thamarailingam Athilingam, Prabhat Tiwari, Prachi Pathak, Shaobo Zhang, Sophie Theis, Timothy E Saunders

**Affiliations:** Warwick Medical School and Centre for Mechanochemical Cell Biology, University of Warwick, Gibbet Hill Road, Coventry, CV4 7AL, United Kingdom; Mechanobiology InsItute, NaIonal University of Singapore, 117411, Singapore

## Abstract

The use of fluorescent proteins has become ubiquitous throughout the life sciences as a key method for understanding molecular, cellular and tissue scale processes. Genetically encodable fluorophores have enabled stable genetic lines to be generated in a large array of organisms. There are now suites of fluorophores available, particularly in the green and red spectra. Yet, which fluorophore works best *in vivo* can depend on a range of factors, both extrinsic (*e*.*g*. pH, temperature) and intrinsic (*e*.*g*. photobleaching, brightness). While fluorophores have been well characterised in cell culture, such measures within *in vivo* systems are more limited. Here, we present a quantitative screen of nine green and eight red fluorophore lines in *Drosophila*, with the fluorescent protein expressed from the same genomic location and imaged under identical conditions. We analyse the expression of the fluorophores in both early and late *Drosophila* embryos. We provide a quantitative analysis of the bleaching and folding rates. We find amongst the green fluorophores that the suitable choice – *e*.*g*., mEGFP, mNeonGreen, mStayGold - depends on timing and imaging requirements. For the red fluorophores, mScarlet-I performed consistently well, though no particular fluorophore stood out as ideal under all conditions. These results provide a powerful database for selecting optimal fluorophores for imaging in the *Drosophila* embryo in green and red channels.

## Introduction

Genetically encodable fluorescent proteins have become a central tool in much of developmental biology^1–3^. Quantification of spatiotemporal dynamics in genetic processes and morphogenesis has led to breakthroughs that would have not been possible in fixed samples^4^. Since the original isolation and use of GFP^5–7^, there has been a huge expansion in the available fluorophore lines, taking advantage of targeted mutagenesis^8^ and also fluorophores from different species^9,10^. This has included the generation of fluorophores activatable at longer wavelengths, enhancing *in vivo* imaging.

Many factors determine the efficacy of specific fluorophores, including brightness, folding time, sensitivity to pH, photo-bleaching and -toxicity, and stability^11–15^. There has been continual improvement in the claimed efficacy of fluorophores, exemplified by the recent work on StayGold, an especially photostable green fluorophore^16–19^, and variants of mScarlet that display fast maturation for a red fluorophore^20^. Testing of fluorophores has typically been in cell culture^21–23^ or relatively simple systems such as yeast^24,25^ or *C elegans*^26–28^. Yet, fluorophore behaviour varies between systems and even developmental stages^29^. In Zebrafish and Medaka, there has been comparison of fluorophore intensity^29^ but no rigorous exploration of folding time or bleaching *in vivo*. To make an informed decision about the right fluorophore to use for a specific experiment requires knowledge of its intensity, folding rate and stability in the organism of interest. It is also important to minimise experimental variation; *e*.*g*., ensuring that each fluorophore is expressed from the same genomic location.

Here, we analyse popularly used monomeric green and red fluorophores in *Drosophila* (Fig. S1), derived from different species (Fig. S2). We took advantage of *Drosophila* genetics to ensure all insertions are at the same genomic position. We utilised the same spinning disc confocal microscope (Methods) for all experiments, due to the ubiquity of such systems in the life sciences for *in vivo* imaging. We report brightness in both early and late embryo stages, as well as folding and bleaching rates. With these results, researchers can select appropriate fluorophores for specific experiments in the *Drosophila* embryo.

## Results

### Generation of fluorescent reporter lines

We opted for fluorophores reported in other systems and from FPbase as bright, fast maturing, highly stable, and monomeric forms wherever possible, with only superfolder GFP displaying a weak dimerisation (Fig. S1 and Fig. S2). We generated stable lines expressing histone H2Av-fluorophore fusions inserted at the same genomic location (Fig. S3), as described in Methods. For each insertion we generated around 3-6 independent sub-lines. Each sub-line was checked for consistent expression (Fig. S4). We note that most sub-lines were very similar in behaviour but some displayed anomalous fluorescence and these were discarded first. From the sub-lines with similar intensity and fluorescence distribution, we maintained a single line per insertion to minimise husbandry time. Each line was homozygous viable and showed no clear phenotype.

### Fluorescence intensity in the early embryo

We first assayed the fluorescence intensity in the early *Drosophila* embryo. The blastoderm stage of *Drosophila* development is highly amenable for live imaging^30–35^. The observed intensity depends on the molecular brightness of the fluorophore and the amount of folded protein. Bright proteins may display a low fluorescence intensity in the early embryo due to a relatively low fraction of folded protein.

We carefully staged embryos in nuclear cycle (n.c.) 14, 20 ± 2 mins post the 13^th^ nuclei division. Embryos were mounted dorsally and z-stacks were acquired using a spinning disc confocal microscope (Fig. 1A-B, Methods). The fluorophores have different reported folding times, resulting in a broad range of observed intensity. We segmented each nucleus (Methods) to ensure we only described the nuclear signal. As we were comparing embryos with the same genetic insertion location, recorded on the same microscope under identical imaging and environmental conditions we did not perform background subtraction on our intensity measurements. This was to negate any potential artefacts from the background subtraction algorithm selected. Quantifying the measured intensity across at least three embryos for each fluorescent reporter, we saw that in the early embryo mEmerald and mScarlet-I resulted in the highest intensity signal for green and red fluorophores, respectively (Fig. 1C-D, Fig. S5A-B, Movie S1).

**Figure 1:**
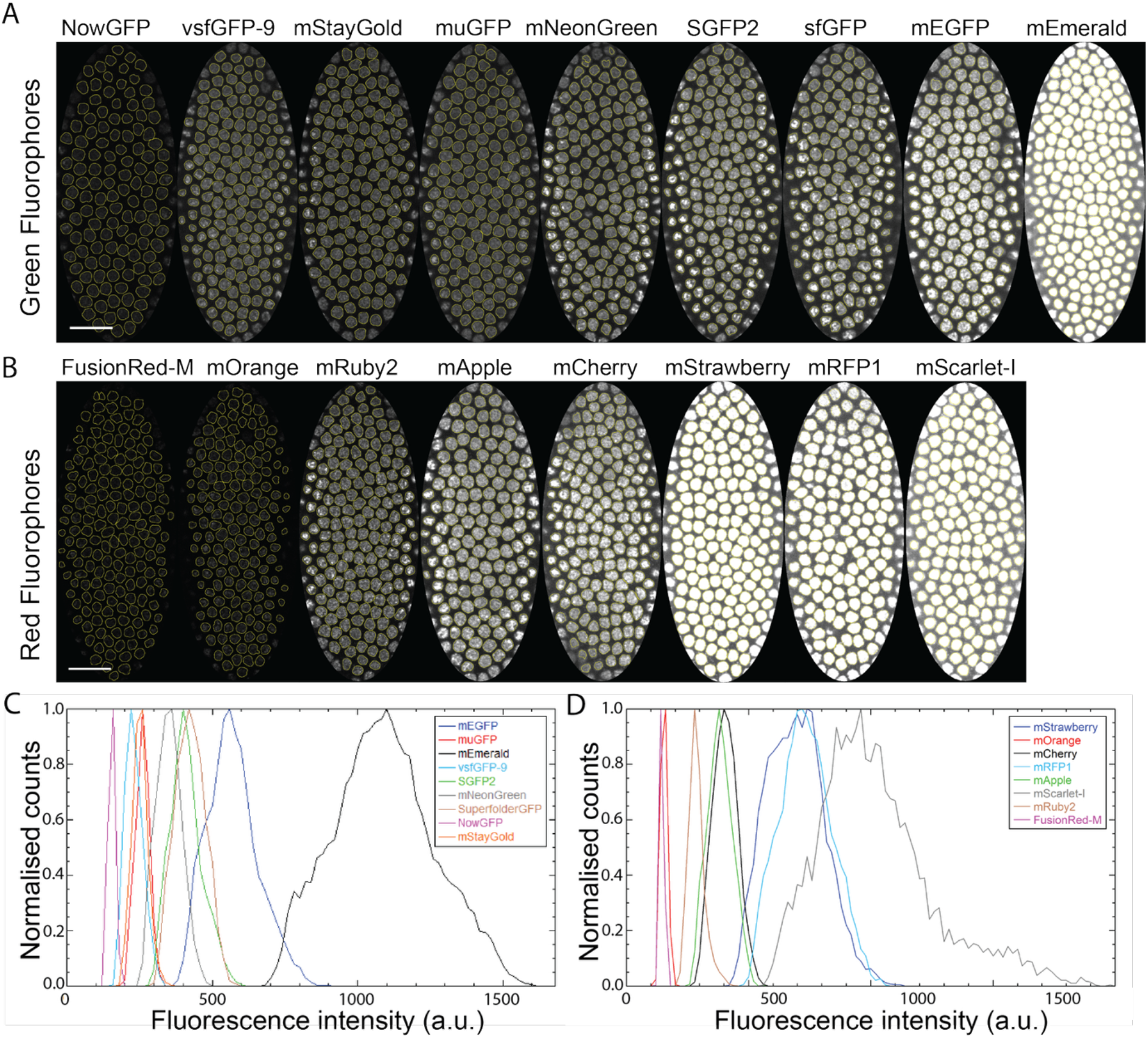
Fluorophore intensity in the *Drosophila* blastoderm (n.c. 14). Comparison of (**A**) green and (**B**) red fluorescence intensity using the same intensity scaling in n.c. 14. The fluorescence signal did not saturate. Shown are single imaging planes. Normalised histograms of fluorescence intensity for green (**C**) and red (**D**) fluorophores averaged across at least n=3 embryos per line. Scale bars = 20 μm.

### Fluorescence intensity during embryo stage 15-16

We next looked at stage 15-16 embryos, kept at room temperature (22ºC). This stage corresponds to the end of germband retraction and completion of internal organ formation, such as the heart, nerve cord and gut. Embryos were mounted dorsally and we collected a z-stack through each embryo (Methods).

We imaged embryos at the stage of heart closure and ordered them by intensity (Fig. 2A-B). Again, we see a clear hierarchy in the intensity, though with some changes in order compared to the early embryo. For example, mEGFP performs best amongst the green proteins. mStayGold is relatively more intense than in the early embryo, though it is still comparatively less fluorescent than a number of other GFPs. In the red proteins, mStrawberry performed best later, with mScarlet-I comparatively less intense. These results are summarised in Fig. 2C-D and Fig. S5C-D.

**Figure 2:**
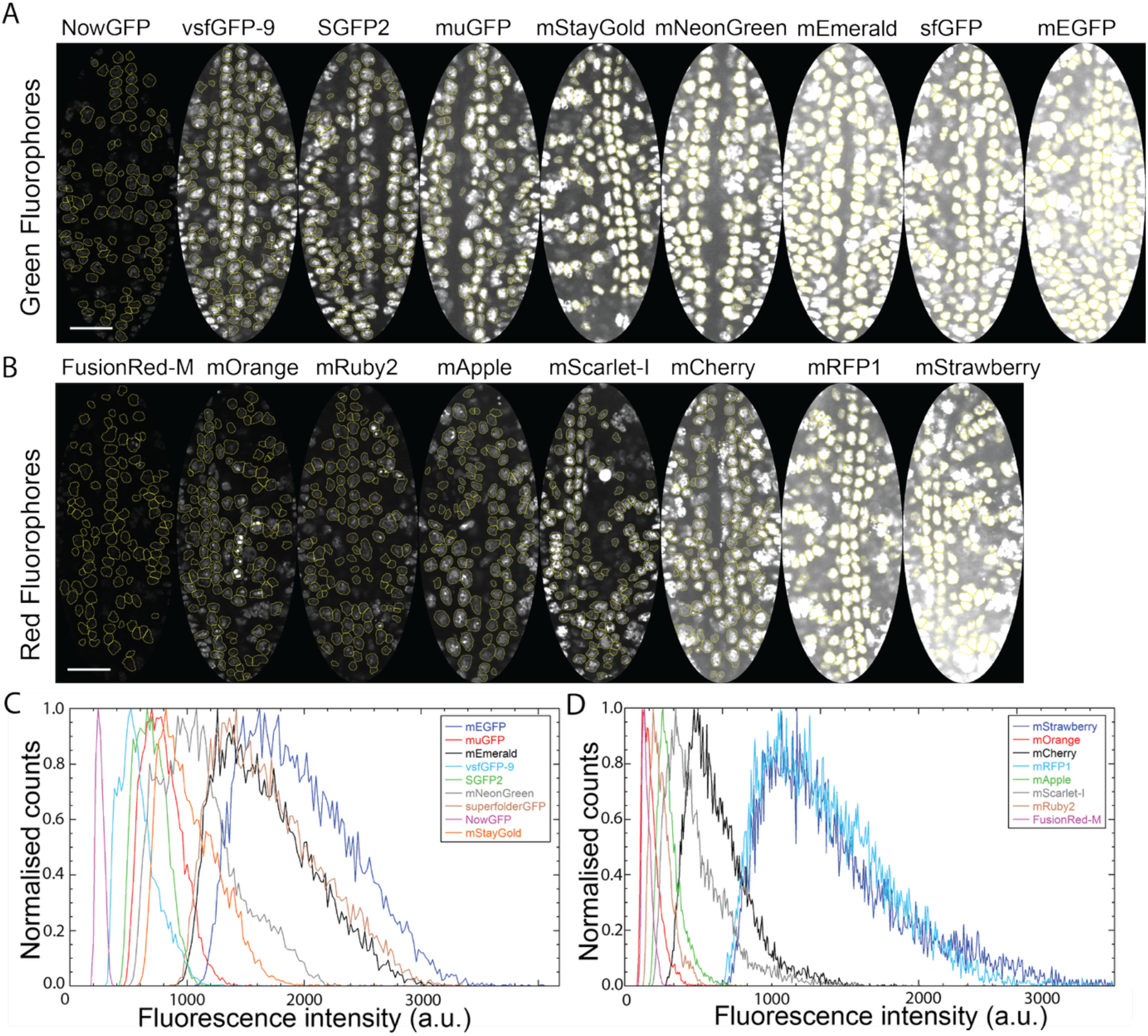
Fluorophore intensity in the *Drosophila* stage 15 embryo. Comparison of (**A**) green and (**B**) red fluorescence intensity using the same intensity scaling at stage 15. Shown are single imaging planes. The fluorescence signal did not saturate. Normalised histograms of fluorescence intensity for green (**C**) and red (**D**) fluorophores. Each curve corresponds to at least n=3 embryos. Scale bars = 20 μm.

### Fluorophore folding time

The fluorescence intensity is not the only important factor in choosing a fluorophore. For example, the folding time and the stability are also necessary for consideration. We next focused on the fluorophore folding time.

Males of each of the green or red fluorescent protein expressing lines were crossed with females expressing Nup107, which marks the nuclear envelope. For our green fluorescent lines we crossed with Nup107::mRFP and for the red fluorescent lines we crossed with Nup107::mEGFP (Methods). tince the maternal-zygotic transition occurred the fluorophores were transcribed, leading to increased fluorescence through development (Fig. 3A-B, Fig. S6, Movie S2). We analysed the increase in intensity over time for both green and red fluorophores. Time point zero was defined as onset of gastrulation for both the green and red fluorescent proteins. Given no signal was expected at experiment onset, we subtracted background such that the initial signal was close to zero. The green lines all showed similar behaviour, with an increase in signal above background first appearing around 120 minutes post imaging start (Fig. 3C). Estimation of the fluorophore folding time is difficult as the intensity profiles are functions of H2Av mRNA production and turnover, protein folding rate, and protein degradation. Further, there can be changes in signal due to morphological changes in the embryo; *e*.*g*., gastrulation and germband elongation change the cell position. It was apparent that by the end of the imaging, the green fluorescence signal had not reached a steady-state for most fluorophores (Fig. 3C). Indeed, we observed increases in signal that may correspond to changes in the fluorophore production rate during the experiment.

**Figure 3:**
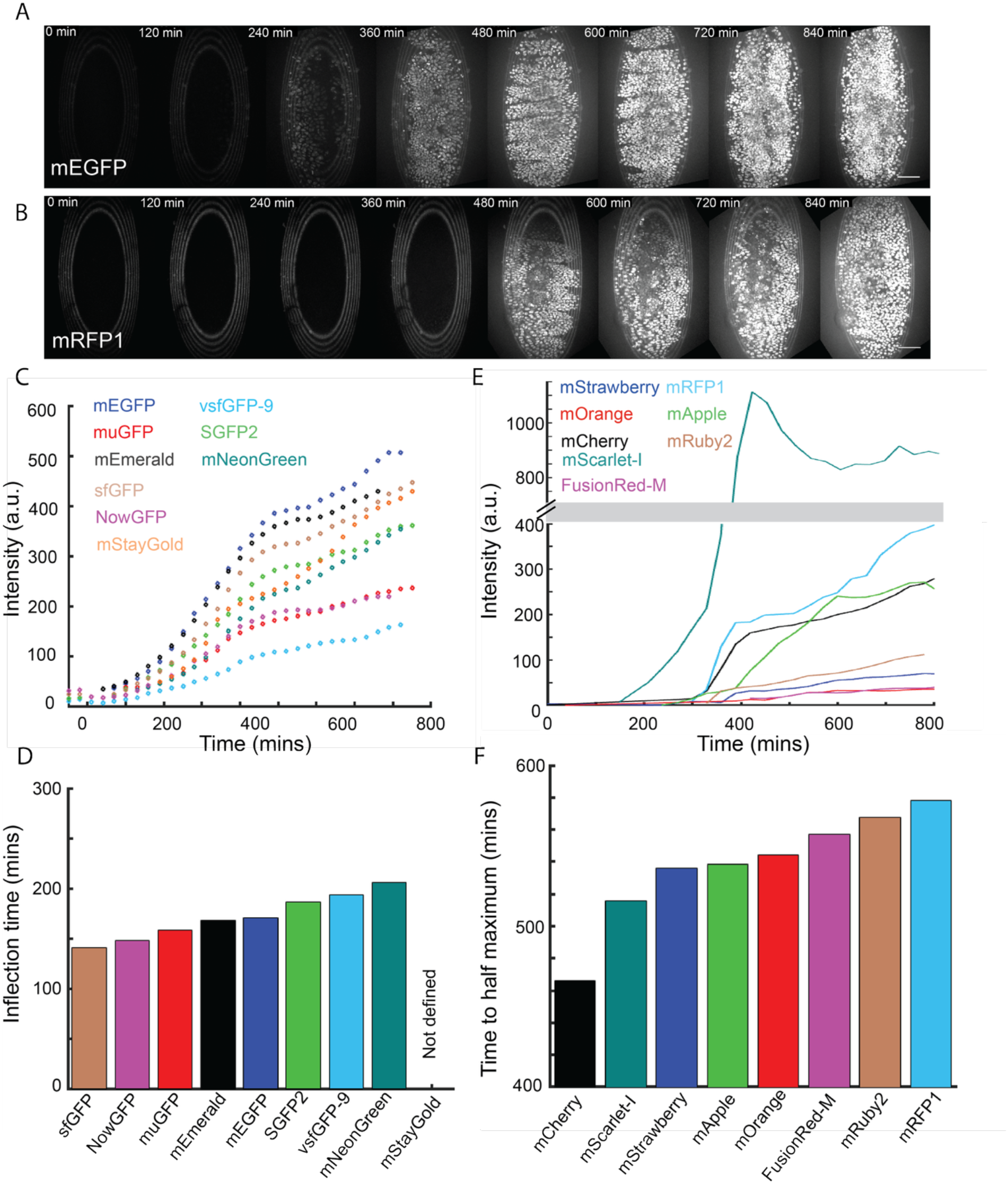
Fluorophore folding during *Drosophila* embryogenesis. (**A-B**)) mEGFP and mRFP1 fluorescence intensity across time, driven only by zygotic transcription. Shown are maximum intensity projections (scale bars = 30 μm). (**C**) Intensity of green fluorescent proteins over time. Time 0 corresponds to onset of gastrulation. (**D**) Time to inflection point in green fluorescent protein intensity profiles (Methods). (**E**) Intensity of red fluorescent proteins over time. Time 0 as in (C). (**F**) Time for red fluorescent proteins to reach half maximum intensity. At least n=3 embryos for each fluorophore.

Therefore, to estimate the folding rate of the green fluorescent proteins we looked at when the fluorescent profiles reached the inflection point (*t*_*in*_), defined by the phenomenological 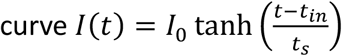 (Fig. 4D, Methods). *t*_*s*_ is related to the time to reach steady-state, with larger *t*_*s*_ denoting larger time to obtain a steady-state. We see that the inflection time is smallest for Superfolder GFP and longest for mNeonGreen (Fig. 4D). The only profile that did not fit well to this phenomenological function was mStayGold, which did not display a clear reduction in its rate of intensity increase. From examining the curves in Fig. 3C, we see that mStayGold expression appears later than most of the fluorophores. A key point from Fig. 3C is that the difference in timing between the different green fluorophores was small. Indeed, normalising by the final intensity and applying a time offset, the folding curves for the green fluorescent proteins largely fell upon each other (Fig. S7A). This suggests that the dynamic behaviour of folding was similar across the green fluorophores.

**Figure 4:**
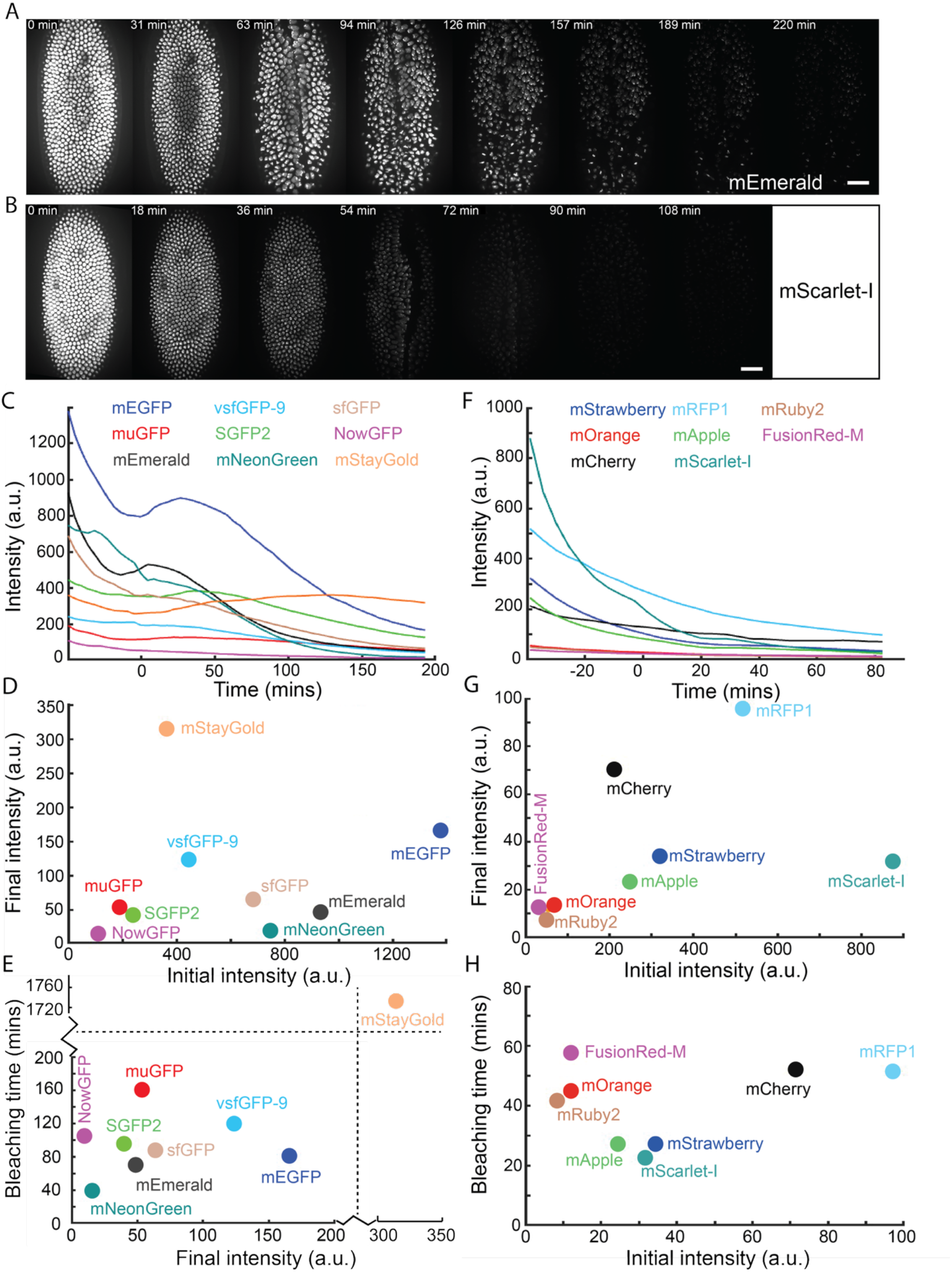
Fluorophore bleaching in the *Drosophila* embryo. (**A-B**) mEmerald and mScarlet-I fluorescence intensity across time, with continued illumination over 4 and 2 hours, respectively (Methods). Scale bars = 30 μm. (**C**) Intensity of green fluorescent proteins (FPs) over time. Time 0 corresponds to onset of gastrulation. (**D**) Comparison of initial to final (4 hours later) intensities for the green FPs. (**E**) Comparison of final intensity to the estimated bleaching time for green FPs. (**F**) Intensity of red FPs over time, Time 0 as in (C). (**G**) As (**D**) but for the red FPs (illumination for 2 hours). (**H**) Comparison of initial intensity to the estimated bleaching time for red FPs. At least n=3 embryos for each fluorophore.

For the red fluorophores, the intensity profiles were more varied (Fig. 3E). They could not be reduced to a single curve, unlike the green fluorophores (Fig. S7B). The phenomenological function *I*(*t*) did not fit the curves well. Therefore, to estimate the folding time, we calculated: (i) the time to reach half maximum intensity; and (ii) the time for the signal to reach 25 a.u. (all embryos imaged under similar conditions on the same microscope). We see that mCherry and mScarlet reach their half maximum intensity comparatively quickly, whereas mRFP1 and mRuby2 are over an hour slower (Fig. 3F). The initiation time (Fig. S7C) showed similar trends between the fluorophores.

To conclude, amongst the green fluorophores analysed, there were no major differences in apparent folding rate. All fluorophores started to increase around 100-150 minutes post gastrulation. In contrast, the red fluorophores displayed more varied intensity profiles, with large differences in both when signal appeared and the intensity obtained during the experiment. tiverall, mCherry and mScarlet-I performed well considering onset of signal and subsequent intensity.

### Fluorophore stability

Along with the folding time, another important factor in selecting a fluorophore is protein stability. We imaged each line with the same laser intensity over the same developmental period. tif course, during this period new fluorophores are generated so we are not measuring the photobleaching rate alone. However, this measurement mirrors experimental situations and provides a useful benchmark for comparing fluorophore performance *in vivo*.

We started imaging ventrally mounted embryos during n.c. 14, with time zero defined as the onset of gastrulation. We recorded f or four and two hours for the green and red embryos respectively (Methods). Example embryos are shown in Fig. 4A-B and Movie S3-S4 and across all fluorophore lines in Fig. S8A-B. We used 21.8mW and 38.5mW laser power for the green and red fluorophores respectively. We imaged with 400ms exposure time and every 4.5 minutes for both red and green fluorophores. With these conditions we saw clear bleaching curves, with a small number of embryos appearing nonviable towards the end of imaging for the green fluorophores, which were excluded from the analysis. Accurately measuring the photobleaching rate is very challenging, as it is an amalgamation of different factors^14,22,36,37^ (see discussion at https://www.fpbase.org/bleaching/). Bleaching rates do not scale linearly with laser power^22^. We imaged our embryos under the same conditions, with laser power and imaging rates chosen so that a significant majority of the embryos were apparently healthy at the end of imaging.

For the green fluorescent proteins, we observed non-monotonic behaviour in the intensity profiles (Fig. 4C). Fluorophores including mEGFP and mEmerald displayed a secondary peak in intensity around an hour aler experiment initiation. This is consistent with a change in the rate of protein production. Despite this, we can follow overall trends in the fluorophor e behaviour. We fitted the phenomenological function 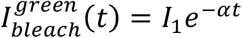 to the bleaching curve s, with α approximating the bleaching time. For the fluorophores displaying a secondary peak in intensity, we fitted this function to the time points aler this peak (Fig. S8C). mEGFP is very bright but undergoes significant bleaching (Fig. 4D-E). Due to its high initial intensity, mEGFP is still more intense aler four hours than other fluorophores that showed a less severe bleaching response. vsfGFP-9 performed strongly, with slow bleaching. muGFP also had slow bleaching but concomitantly low intensity.

Consistent with previous reports^16–18^, mStayGold displayed remarkable photostability. In terms of initial intensity, five other green fluorophores had higher signal at the beginning of the experiment. Four hours later, mStayGold maintained its intensity level (in part due to new protein expression), with a signal nearly twice that of the next brightest fluorophore. Therefore, for long-term imaging in the *Drosophila* embryo, mStayGold appears to be an excellent choice.

The bleaching experiment for the red fluorescent proteins displayed a more monotonic behaviour (Fig. 4F). All bleaching curves were approximated well by the formula 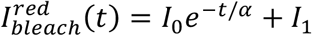, where *I*_0,1-_ and α are fimng constants. In particular, α provides an approximation of the bleaching time of the fluorophore. We emphasise that this is a phenomenological model, and incorporates multiple potential factors that impact the parameters^14,22,36,37^.

In our bleaching analysis, mRFP1 performed strongly, with high intensity (Fig. 4G) and one of the longest bleaching times (Fig. 4H). FusionRed-M bleached slowly but had low intensity. mScarlet-I had brightest initial intensity but had the shortest phenomenological bleaching time. We conclude that for long-term imaging in the *Drosophila* embryo mRFP1 and mCherry are good options.

## Discussion

We have carefully compared the efficacy of 17 fluorophores within the *Drosophila* embryo. We have considered early and late stages of development, which has revealed that the most suitable fluorophore is highly dependent on the imaging requirements. mStayGold is remarkably stable as reported previously, but is not as bright within the embryo as other fluorophores. For short imaging or low expressing proteins, other green fluorophores may still be more applicable. We saw a large variation within the red family of fluorophores.

These proteins typically had greater variation in their folding time than the green fluorophores, which impacts their use *in vivo*, especially early in development. mScarlet-I consistently performed well though it bleached relatively quickly (somewhat compensated for by its high intensity).

While we have focused on genetically encodable fluorescent proteins in the green and red spectra, there are a broad range of fluorophores available, from blue to far red^38^. It will be interesting to perform a similar analysis on these. Far-red fluorophores have been assayed in cell culture and *C elegans*^39^. Further, there are now many fluorescent dyes, many of which are compatible with live imaging *in vivo*. There has been rigorous analysis of fluorescent dyes in fixed systems^40^, but as far as we are aware, not in complex living systems. tiur results here demonstrate that extrapolating from *in vitro* or cell culture systems to *Drosophila* can lead to misleading choices of fluorophores.

There are further image analysis tools that may alter our interpretations. We did not deconvolve our images^41^. There are also machine-learning denoising approaches^42,43^ that could, potentially, be more efficient with different fluorophores. As machine-driven approaches increase in power, *e*.*g*., segmentation^44,45^, how the data from the microscope aligns with these techniques may impact the final quality of the quantitative data extraction. Finally, we have not performed cell tracking here. It would be interesting to test how different fluorophores perform with cell tracking algorithms^46^.

## Acknowledgements

We thank the Mechanobiology Institute and Warwick Medical School for support through seed and startup grants respectively, and provision of fly rooms. We thank the *Drosophila* community for feedback on the approach. We thank Mohan Balasubramanian for sharing mStayGold.

## Author Contributions

TES designed and managed the study. BCYL collected and analysed data with ST. TA and PT generated the constructs for the fly lines. TA confirmed the insertions. PD, PT and SZ collected initial data and helped design the study. BYCL, TA, PD and TES carried out fly maintenance and selection. BCYL and TES generated the figures and movies. TES and BCYL wrote the first dral of the manuscript with all authors contributing to the final submission.

## Methods

### Generation of Histone (H2Av) fused fluorophore lines

We used the PhiC31 mediated recombination method to generate stable transgenic lines of Histone H2Av fused fluorescence proteins (FPs). Initially, the Histone H2Av gene sequence, including the 5’ and 3’ UTRs was cloned into XhoI and BamHI sites of w+ajB plasmid (Addgene 30326). The restriction sites, HindIII, BglII, and NotI were engineered in tandem before the stop codon of H2Av C’ end. The gene fragments of each of the FPs were codon optimized manually for their stable expression in *Drosophila* embryos (Supplementary Table 1 and 2). The FP gene fragments with the necessary codon changes flanked with BglII and NotI sites at their N and C termini, respectively, were inserted at H2Av C’-end aler restriction digestion, just before the H2Av 3’UTR. A flexible (GS)_3_ linker was inserted between the HindIII and BglII sites at the N-terminus of each FP sequence.

The H2Av::FP transgenic lines were generated by PhiC31 mediated recombination by injecting the H2Av::FP-w+ajB transgene plasmid into the ajP2 (chro.III) landing site acceptor line BL8622. The insertions were carried out by BestGene U.S.A. The multiple lines of each of the fluorescent transgenic lines were checked for any variations in brightness.

The insertion sites of all the FP transgenic lines were validated by PCR and sequencing (Fig. S3).

### Sample preparation

#### Nuclear cycle (n.c.) 14 sample preparation

Flies were placed in apple-juice agar sealed cages and fed with yeast paste for two days before eggs were collected. Aler 1 hour of collection, the eggs were kept at room temperature (~20ºC) for two hours and then dechorionated by incubating them in 50% bleach for 3 minutes. tince rinsed and recovered on a mesh aler bleaching, the embryos were mounted dorsally on Fluorodishes (FD35, World Precision Instruments) using Heptane glue and covered with a drop of Halocarbon oil 700 (H8898, SIGMA-ALDRICH). The embryos were transferred to the microscope stage and imaged 20 minutes aler the last syncytial nuclear division (n.c. 14). The photobleaching experiment preparation was similar. In the case of the protein folding experiments, the average start of imaging was 50 ± 18 mins and 61 ± 20 mins prior to gastrulation for the green and red fluorophores respectively. For the photobleaching experiments, the average start of imaging was 38±15 mins and 39±23 mins prior to gastrulation for the green and red fluorophores respectively.

#### Stage 15-16 sample preparation

Flies were placed in apple-juice agar sealed cages and fed with yeast paste for two days before eggs were collected. Aler 2 hours of collection, the eggs were kept at room temperature (~20ºC) for ~ 15 hours (overnight) before being processed like the early-stage embryos. The embryos were imaged at stage 15-16, around the time of heart closure.

#### Embryo preparation for maturation experiments

Males of each of the green or red fluorescent protein expressing lines were crossed with females of line BL 35517 (w*; wgSp-1/Cyti; P(mRFP-Nup107.K)7.1) or line BL 35513 (w*, P(GFP-Nup107.K)13.2.1; wgSp-1/Cyti), respectively. The crosses were kept at room temperature (~20ºC) in a vial with food for one day, and then the flies were placed in apple-juice agar sealed cages with yeast paste for an additional two days, aler which, the embryo samples were prepared as already described above (n.c. 14 sample preparation).

### Imaging conditions

Imaging was performed predominantly in stage 4-5 (early) or stage 15-16 of *Drosophila* development. Embryos were mounted dorsally for data collected in Figs. 1–2. For the bleaching and folding experiments, embryos were mounted ventrally. tinly embryos mounted in the same orientation were compared for statistical analysis. Imaging was performed at 22±1ºC for Figures 1–2 and at 25±1ºC for Figure 3–4. All imaging was performed on an Evident Spinning Disc Confocal Microscope. We recognise that spinning disc microscopy introduces potential sources of error (cross-talk between apertures, type of disc used^47,48^), but it represents a broadly used and accessible system across the life sciences for live imaging live samples. With recent improvements in technology, spinning disc microscopy can generate quantitative data, such as single molecule counting^49^.

### Imaging details

| Experiment | Z-slices | Z-step (um) | Time interval (min) | Exposure time (ms) | Laser power (mW) |
|---|---|---|---|---|---|
| Early green (stage 5) | 51 | 0.5 | NA | 10<sup>3</sup> | 3.9 |
| Late green (stage 15-16) | 100 | 0.5 | NA | 10<sup>3</sup> | 3.9 |
| Green bleaching | 81 | 0.5 | 4.5 | 400 | 21.8 |
| Green maturation | 11 | 2 | 30 | 10<sup>3</sup> | 3.9 |
| Early red | 51 | 0.5 | NA | 10<sup>3</sup> | 38.5 |
| Late red | 100 | 0.5 | NA | 10<sup>3</sup> | 38.5 |
| Red bleaching | 81 | 0.5 | 4.5 | 400 | 38.5 |
| Red maturation | 11 | 2 | 30 | 10<sup>3</sup> | 38.5 |

60X (NA=1.3) silicone oil objective used in all experiments. Approach motivated by similar experiments in Ref. <sup>50</sup>.

### Image analysis

Image analysis was done using custom Fiji macros. The macros are listed below together with a description of what they were used for. The Macros are available on hjps://github.com/TimSaundersLab

Briefly, the “stage analysis macros” produce the following files for each embryo analysed:

1. Ellip’cal RtiI where segmenta’on and measurements were made.
2. Image of the distribu’on of the fluorescence measurements.
3. Table of all the individual fluorescence measurements, in addi’on to measurements of addi’onal parameters.
4. RtiIs where the individual measurements were made.

We analysed the slices from 7μm to 20.5μm depth.

The results corresponding to each embryo of each fly line were combined to generate a single fluorescence intensity distribution, and all fly lines distributions were then plojed together to be compared. Figures were generated using Matlab.

### Fluorophore maturation and bleaching

The maturation and bleaching analysis macros produce the following files for each embryo analysed:

1. Ellip’cal RtiI where segmenta’on and measurements were made.
2. Table with mean fluorescence, standard devia’on, and number of segmented objects, per frame in all cases.
3. Plot of the mean fluorescence per frame, including standard devia’on.
4. Table of all the individual fluorescence measurements, in addi’on to measurements of addi’onal parameters.
5. RtiIs where the individual measurements were made.
6. Table with summarised results per slice/frame.

Depth 9 μm to 21 μm was analysed. Imaging started 7 μm deep into the embryos instead of at the site where the embryos touch the glass onto which they were mounted, which is the case for all other experiments.

The algorithm used to segment the red nuclei was different to that used for the segmentation of the green nuclei. We adapted each one to account for the different fluorescence levels; especially the low intensity in some red lines. In the lowest frames, there were false positives which were removed manually.

The macros used for the analysis of the fluorescence in the maturation experiments include a step to input the frame to be considered as time point 0. This time synchronisation was performed by visual inspection of the images of the embryos, focusing on the onset of gastrulation when possible.

For each group *i*, we calculated the number of elements *n*_2_, the standard deviation *σ*_2_ and the mean 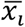. We calculated the combined mean as follows:

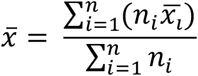

The combined stan dard deviation was then calculated according to:

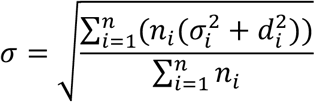

where *d*_2_ is the difference between the combined mean 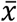 and the mean of the group 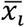.

### Model Fiting

#### Folding rates

We fitted the phenomenological function 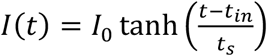 to the green fluorescent protein folding data. We used the function *fit* in Matlab to fit the parameters. For the red fluorophores, *I*(*t*) only described well a subset of the curves. Therefore, we used the time to half maximum to infer knowledge of the folding rate.

#### Bleaching rates

We fitted the function 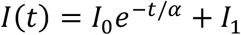 to the red and green profiles using the *fit* function in Matlab. For the green proteins, we only fitted the curves after the intermediate increase in signal (Fig. S8C).

