## Supplementary Material for "Identification of optimal fluorophores for use in the *Drosophila* embryo"

**Contents:**

1. **Supplementary Tables**
2. **Supplementary Figures**
3. **Supplementary Video Legends**
4. **Supplementary Analysis of Folding Rates**
5. **Supplementary References**

### Supplementary Table 1

| Green Fluorophores | Codon optimized Gene Sequence |
|---|---|
| mEmerald | AGATCTGTGAGCAAGGGCGAGGAGCTGTTCACCGGGGTGGTGCCCATCCTGGTCGAGCTGGAC<br>GGCGACGTAAACGGCCACAAGTTCAGCGTGTCCGGCGAGGGCGAGGGCGATGCCACCTACGG<br>CAAGCTGACCCTGAAGTTCATCTGCACCACCGGCAAGCTGCCCGTGCCCTGGCCCACCCTCGT<br>GACCACCTTGACCTACGGCGTGCAGTGCTTCGCCCGCTACCCCGACCACATGAAGCAGCACGA<br>CTTCTTCAAGTCCGCCATGCCCGAAGGCTACGTCCAGGAGCGCACCATCTTCTTCAAGGACGAC<br>GGCAACTACAAGACCCGCGCCGAGGTGAAGTTCGAGGGCGACACCCTGGTGAACCGCATCGAG<br>CTGAAGGGCATCGACTTCAAGGAGGACGGCAACATCCTGGGGCACAAGCTGGAGTACAACTACA<br>ACAGCCACAAGGTCTATATCACCGCCGACAAGCAGAAGAACGGCATCAAGGTGAACTTCAAGAC<br>CCGCCACAACATCGAGGACGGCAGCGTGCAGCTCGCCGACCACTACCAGCAGAACACCCCCAT<br>CGGCGACGGCCCCGTGCTGCTGCCCGACAACCACTACCTGAGCACCCAGTCCAAGCTGAGCAA<br>AGACCCCAACGAGAAGCGCGATCACATGGTCCTGCTGGAGTTCGTGACCGCCGCCGGGATCAC<br>TCTCGGCATGGACGAGCTGTACAAG**TAA** |
| mEGFP | AGATCTGTGAGCAAGGGCGAGGAGCTGTTCACCGGGGTGGTGCCCATCCTGGTCGAGCTGGAC<br>GGCGACGTAAACGGCCACAAGTTCAGCGTGTCCGGCGAGGGCGAGGGCGATGCCACCTACGG<br>CAAGCTGACCCTGAAGTTCATCTGCACCACCGGCAAGCTGCCCGTGCCCTGGCCCACCCTCGT<br>GACCACCCTGACCTACGGCGTGCAGTGCTTCAGCCGCTACCCCGACCACATGAAGCAGCACGA<br>CTTCTTCAAGTCCGCCATGCCCGAAGGCTACGTCCAGGAGCGCACCATCTTCTTCAAGGACGAC<br>GGCAACTACAAGACCCGCGCCGAGGTGAAGTTCGAGGGCGACACCCTGGTGAACCGCATCGAG<br>CTGAAGGGCATCGACTTCAAGGAGGACGGCAACATCCTGGGGCACAAGCTGGAGTACAACTACA<br>ACAGCCACAACGTCTATATCATGGCCGACAAGCAGAAGAACGGCATCAAGGTGAACTTCAAGATC<br>CGCCACAACATCGAGGACGGCAGCGTGCAGCTCGCCGACCACTACCAGCAGAACACCCCCATC<br>GGCGACGGCCCCGTGCTGCTGCCCGACAACCACTACCTGAGCACCCAGTCCAAGCTGAGCAAA<br>GACCCCAACGAGAAGCGCGATCACATGGTCCTGCTGGAGTTCGTGACCGCCGCCGGGATCACT<br>CTCGGCATGGACGAGCTGTACAAG**TAA** |
| Superfolder GFP | AGATCTAGCAAGGGCGAGGAGCTGTTCACCGGGGTGGTGCCCATCCTGGTCGAGCTGGACGGC<br>GACGTAAACGGCCACAAGTTCAGCGTGCGCGGCGAGGGCGAGGGCGATGCCACCAACGGCAA<br>GCTGACCCTGAAGTTCATCTGCACCACCGGCAAGCTGCCCGTGCCCTGGCCCACCCTCGTGAC<br>CACCCTGACCTACGGCGTGCAGTGCTTCAGCCGCTACCCCGACCACATGAAGCGCCACGACTTC<br>TTCAAGTCCGCCATGCCCGAAGGCTACGTCCAGGAGCGCACCATCAGCTTCAAGGACGACGGC<br>ACCTACAAGACCCGCGCCGAGGTGAAGTTCGAGGGCGACACCCTGGTGAACCGCATCGAGCTG<br>AAGGGCATCGACTTCAAGGAGGACGGCAACATCCTGGGGCACAAGCTGGAGTACAACTTCAACA<br>GCCACAACGTCTATATCACCGCCGACAAGCAGAAGAACGGCATCAAGGCCAACTTCAAGATCCG<br>CCACAACGTGGAGGACGGCAGCGTGCAGCTCGCCGACCACTACCAGCAGAACACCCCCATCGG<br>CGACGGCCCCGTGCTGCTGCCCGACAACCACTACCTGAGCACCCAGTCCGTGCTGAGCAAAGA<br>CCCCAACGAGAAGCGCGATCACATGGTCCTGCTGGAGTTCGTGACCGCCGCCGGGATCACTCA<br>CGGCATGGACGAGCTGTACAAG**TAA** |
| SGFP2 | AGATCTGTGAGCAAGGGCGAGGAGCTGTTCACCGGGGTGGTGCCCATCCTGGTCGAGCTGGAC<br>GGCGACGTAAACGGCCACAAGTTCAGCGTGTCCGGCGAGGGCGAGGGCGATGCCACCTACGG<br>CAAGCTGACCCTGAAGTTCATCTGCACCACCGGCAAGCTGCCCGTGCCCTGGCCCACCCTCGT<br>GACCACCCTGACCTACGGCGTGCAGTGCTTCGCCCGCTACCCCGACCACATGAAGCAGCACGA<br>CTTCTTCAAGTCCGCCATGCCCGAAGGCTACGTCCAGGAGCGCACCATCTTCTTCAAGGACGAC<br>GGCAACTACAAGACCCGCGCCGAGGTGAAGTTCGAGGGCGACACCCTGGTGAACCGCATCGAG<br>CTGAAGGGCATCGACTTCAAGGAGGACGGCAACATCCTGGGGCACAAGCTGGAGTACAACTACA<br>ACAGCCACAACGTCTATATCACCGCCGACAAGCAGAAGAACGGCATCAAGGCCAACTTCAAGATC<br>CGCCACAACATCGAGGACGGCGGCGTGCAGCTCGCCGACCACTACCAGCAGAACACCCCCATC<br>GGCGACGGCCCCGTGCTGCTGCCCGACAACCACTACCTGAGCACCCAGTCCAAGCTGAGCAAA<br>GACCCCAACGAGAAGCGCGATCACATGGTCCTGCTGGAGTTCGTGACCGCCGCCGGGATCACT<br>CTCGGCATGGACGAGCTGTACAAG**TAA** |
| mNeon-Green | AGATCTGTGAGCAAGGGCGAGGAGGATAACATGGCCTCTCTCCCAGCGACACATGAGTTACACA<br>TCTTTGGCTCCATCAACGGTGTGGACTTTGACATGGTGGGTCAGGGCACCGGCAATCCAAATGAT<br>GGTTATGAGGAGTTAAACCTGAAGTCCACCAAGGGTGACCTCCAGTTCTCCCCCTGGATTCTGGT<br>CCCTCATATCGGGTATGGCTTCCATCAGTACCTGCCCTACCCTGACGGGATGTCGCCTTTCCAGG<br>CCGCCATGGTAGATGGCTCCGGATACCAAGTCCATCGCACAATGCAGTTTGAAGATGGTGCCTCC<br>CTTACTGTTAACTACCGCTACACCTACGAGGGAAGCCACATCAAAGGAGAGGCCCAGGTGAAGG |

| | |
|---|---|
| | GGACTGGTTTCCCTGCTGACGGTCCTGTGATGACCAACTCGCTGACCGCTGCGGACTGGTGCA<br>GGTCGAAGAAGACTTACCCCAACGACAAAACCATCATCAGTACCTTTAAGTGGAGTTACACCACT<br>GGAAATGGCAAGCGCTACCGGAGCACTGCGCGGACCACCTACACCTTTGCCAAGCCAATGGCG<br>GCTAACTATCTGAAGAACCAGCCGATGTACGTGTTCCGTAAGACGGAGCTCAAGCACTCCAAGAC<br>CGAGCTCAACTTCAAGGAGTGGCAAAAGGCCTTTACCGATGTGATGGGCATGGACGAGCTGTAC<br>AAG**TAA** |
| vsfGFP-9 | AGATCTAGCAAAGGAGAAGAACTTTTCACTGGAGTTGTCCCAATTCTTGTTGAATTAGATGGTGAT<br>GTTAATGGGCACAAATTTTCTGTCCGTGGAGAGGGTGAAGGTGATGCTACAAACGGAAAACTCAC<br>CCTTAAATTTATTTGCACTACTGGAAAACTACCTGTTCCGTGGCCAACACTTGTCACTACTCTGAC<br>CTATGGTGTTCAATGCTTTTCCCGTTATCCGGATCACATGAAACGGCATGACTTTTTCAAGAGTGC<br>CATGCCCGAAGGTTATGTACAGGAACGCACTATATCTTTCAAAGATGACGGGACCTACAAGACGC<br>GTGCTGAAGTCAAGTTTGAAGGTGATACCCTTGTTAATCGTATCGAGTTAAAGGGCATTGATTTTAA<br>AGAAGATGGAAACATTCTTGGACACAAACTCGAGTACAACTTTAACTCACACAATGTATACATCAC<br>GGCAGACAAACAAAAGAATGGAATCAAAGCTAACTTCAAAATTCGCCACAACGTTGAAGATGGTT<br>CCGTTCAACTAGCAGACCATTATCAACAAAATACTCCAATTGGCGATGGCCCTGTCCTTTTACCAG<br>ACAACCATTACCTGTCGACACAATCTGTCCTTTCGAAAGATCCCAACGAAAAGCGTGACCACATG<br>GTCCTTCTTGAGTTTGTAACTGCTGCTGGGATTACACATGGCATGGATGAGCTCTACAAAGCCCA<br>GGTGCAGTTGGTCGAATCTGGCGGTGCCCTGGTTCAACCGGGTGGTAGCCTGCGTCTGTCCTG<br>TGCGGCGTCTGGCTTCCCAGTTAATCGTTATAGCATGCGTTGGTACCGTCAAGCGCCTGGCAAG<br>GAGCGTGAGTGGGTCGCGGGTATGAGCTCGGCGGGTGATCGTAGCAGCTACGAAGATTCCGTTA<br>AAGGCCGCTTTACCATCAGCCGTGACGATGCGCGCAATACCGTATACCTGCAAATGAACAGCCTG<br>AAGCCGGAGGACACCGCAGTCTATTACTGCAATGTTAATGTGGGTTTTGAATACTGGGGCCAGGG<br>TACGCAAGTTACCGTTTCC**TAA** |
| NowGFP | AGATCTGTGAGCAAGGGCGAGAAGCTGTTCACCGGGGTGGTGCCCATCCTGGTCGAGCTGGAC<br>GGCGACGTAAACGGCCACAAGTTCAGCGTGTCCGGCGAGGGCGAGGGCGATGCCACCTACGG<br>CAAGATGTCCCTGAAGTTCATCTGCACCACCGGCAAGCTGCCCGTGCCCTGGCCCACCCTCAAA<br>ACCACCCTGACCTGGGGCATGCAGTGCTTCGCCCGCTACCCCGACCACATGAAGCAGCACGAC<br>TTCTTCAAGTCCGCCATGCCCGAAGGCTACGTCCAGGAGCGCACCATCTTCTTCAAGGACGACG<br>GCAACTACAAGACCCGCGCCGAGGTGAAGTTCGAGGGCGACACCCTGGTGAACCGCATCGAGC<br>TGAAGGGCGTCGACTTCAAGGAGGACGGCAACATCCTGGGGCACAAGCTGGAGTACAACGCCA<br>TCAGCGGCAACGCCAATATCACCGCCGACAAGCAGAAGAACGGCATCAAGGCCTACTTCACGAT<br>CCGCCACGACGTCGAGGACGGCAGCGTGCTGCTCGCCGACCACTACCAGCAGAACACCCCCAT<br>CGGCGACGGCCCCGTGCTGCTGCCCGACAACCACTACCTGAGCACCCAGTCCAAGCAGAGCAA<br>AGACCCCAACGAGAAGCGCGATCACATGGTCCTGCTGGAGTTCGTGACCGCCGCCGGGATCCC<br>TCTCGGCGCGGACGAGCTGTACAAG**TAA** |
| muGFP | AGATCTAGCAAGGGCGAGGAGCTGTTCACCGGGGTGGTGCCCATCCTGGTCGAGCTGGACGGC<br>GACGTAAACGGCCACAAGTTCAGCGTGCGCGGCGAGGGCGAGGGCGATGCCACCAACGGCAA<br>GCTGACCCTGAAGTTCATCTGCACCACCGGCAAGCTGCCCGTGCCCTGGCCCACCCTCGTGAC<br>CACCCTGACCTACGGCGTGCTGTGCTTCAGCCGCTACCCCGACCACATGAAGCGCCACGACTTC<br>TTCAAGTCCGCCATGCCCGAAGGCTACGTCCAGGAGCGCACCATCAGCTTCAAGGACGACGGC<br>ACCTACAAGACCCGCGCCGAGGTGAAGTTCGAGGGCGACACCCTGGTGAACCGCATCGAGCTG<br>AAGGGCATCGACTTCAAGGAGGACGGCAACATCCTGGGGCACAAGCTGGAGTACAACTTCAACA<br>GCCACAACGTCTATATCACCGCCGACAAGCAGAAGAACGGCATCAAGGCCTACTTCAAGATCCGC<br>CACAACGTGGAGGACGGCAGCGTGCAGCTCGCCGACCACTACCAGCAGAACACCCCCATCGGC<br>GACGGCCCCGTGCTGCTGCCCGACAACCACTACCTGAGCACCCAGTCCGTGCTGAGCAAAGAC<br>CCCAACGAGAAGCGCGATCACATGGTCCTGCTGGAGGATGTGACCGCCGCCGGGATCACTCAC<br>GGCATGGACGAGCTGTACAAG**TAA** |
| mStayGold | AGATCTGCCAGTACGCCTTTTAAATTCCAGTTGAAAGGTACAATAAATGGCAAATCCTTCACGGTT<br>GAAGGAGAGGGTGAGGGCAACAGCCACGAAGGATCCCATAAAGGCAAGTATGTGTGTACGAGTG<br>GCAAACTCCCTATGTCCTGGGCGGCCCTGGGCACCTCCTTTGGATACGGAATGAAATACTACACT<br>AAATATCCGAGTGGCTTGAAAAATTGGTTTCACGAAGTAATGCCGGAAGGCTTTACATACGATCGA<br>CATATTCAGTATAAGGGTGATGGATCGATACATGCTAAGCATCAACACTTCATGAAAAACGGTACCT<br>ACCACAACATCGTGGAATTTACCGGTCAAGACTTCAAGGAAAACAGCCCCGTACTGACAGGCGAT<br>ATGAACGTCTCGCTGCCAAACGACGTTCAACATATCCCGCGCGACGATGGTGTTGAGTGTCCGG<br>TGACTCTCTTGTATCCCCTGCTCAGTGATAAGAGCAAGTGCGTTGAAGCGCACCAAAATACCATAT<br>GTAAACCCTTGCACAACCAACCCGCGCCAGACGTACCCTATCACTGGATTCGAAAGCAATATACA<br>CAGAGCAAGGACGATACAGAGGAGCGCGACCACATCTGCCAGTCCGAGACTCTCGAAGCACAC<br>TTG**TAA** |

**Supplementary Table 1:** Sequences for the selected green fluorescent proteins.

### Supplementary Table 2

| Red Fluorophores | Codon optimized Gene Sequence |
|---|---|
| mRFP1 | AGATCTGCCTCCTCCGAGGACGTCATCAAGGAGTTCATGCGCTTCAAGGTGCGCATGGAGGGCT<br>CCGTGAACGGCCACGAGTTCGAGATCGAGGGCGAGGGCGAGGGCCGCCCCTACGAGGGCACC<br>CAGACCGCCAAGCTGAAGGTGACCAAGGGCGGCCCCCTGCCCTTCGCCTGGGACATCCTGTCC<br>CCTCAGTTCCAGTACGGCTCCAAGGCCTACGTGAAGCACCCCGCCGACATCCCCGACTACTTGA<br>AGCTGTCCTTCCCCGAGGGCTTCAAGTGGGAGCGCGTGATGAACTTCGAGGACGGCGGCGTGG<br>TGACCGTGACCCAGGACTCCTCCCTGCAGGACGGCGAGTTCATCTACAAGGTGAAGCTGCGCG<br>GCACCAACTTCCCCTCCGACGGCCCCGTAATGCAGAAGAAGACCATGGGCTGGGAGGCCTCCA<br>CCGAGCGGATGTACCCCGAGGACGGCGCCCTGAAGGGCGAGATCAAGATGAGGCTGAAGCTGA<br>AGGACGGCGGCCACTACGACGCCGAGGTCAAGACCACCTACATGGCCAAGAAGCCCGTGCAGC<br>TGCCCGGCGCCTACAAGACCGACATCAAGCTGGACATCACCTCCCACAACGAGGACTACACCAT<br>CGTGGAACAGTACGAGCGCGCCGAGGGCCGCCACTCCACCGGCGCC**TAA** |
| mOrange | AGATCTGTGAGCAAGGGCGAGGAGAATAACATGGCCATCATCAAGGAGTTCATGCGCTTCAAGGT<br>GCGCATGGAGGGCTCCGTGAACGGCCACGAGTTCGAGATCGAGGGCGAGGGCGAGGGCCGCC<br>CCTACGAGGGCTTTCAGACCGCTAAGCTGAAGGTGACCAAGGGTGGCCCCCTGCCCTTCGCCT<br>GGGACATCCTGTCCCCTCAGTTCACCTACGGCTCCAAGGCCTACGTGAAGCACCCCGCCGACAT<br>CCCCGACTACTTCAAGCTGTCCTTCCCCGAGGGCTTCAAGTGGGAGCGCGTGATGAACTTCGAG<br>GACGGCGGCGTGGTGACCGTGACCCAGGACTCCTCCCTGCAGGACGGCGAGTTCATCTACAAG<br>GTGAAGCTGCGCGGCACCAACTTCCCCTCCGACGGCCCCGTAATGCAGAAGAAGACCATGGGC<br>TGGGAGGCCTCCTCCGAGCGGATGTACCCCGAGGACGGCGCCCTGAAGGGCGAGATCAAGATG<br>AGGCTGAAGCTGAAGGACGGCGGCCACTACACCTCCGAGGTCAAGACCACCTACAAGGCCAAG<br>AAGCCCGTGCAGCTGCCCGGCGCCTACATCGTCGGCATCAAGTTGGACATCACCTCCCACAACG<br>AGGACTACACCATCGTGGAACAGTACGAACGCGCCGAGGGCCGCCACTCCACCGGCGGCATGG<br>ACGAGCTGTACAAG**TAA** |
| mStrawberry | AGATCTGTGAGCAAGGGCGAGGAGAATAACATGGCCATCATCAAGGAGTTCATGCGCTTCAAGGT<br>GCGCATGGAGGGCTCCGTGAACGGCCACGAGTTCGAGATCGAGGGCGAGGGCGAGGGCCGCC<br>CCTACGAGGGCACCCAGACCGCCAAGCTGAAGGTGACCAAGGGTGGCCCCCTGCCCTTCGCCT<br>GGGACATCCTAACCCCCAACTTCACCTACGGCTCCAAGGCCTACGTGAAGCACCCCGCCGACAT<br>CCCCGACTACTTGAAGCTGTCCTTCCCCGAGGGCTTCAAGTGGGAGCGCGTGATGAACTTCGAG<br>GACGGCGGCGTGGTGACCGTGACCCAGGACTCCTCCCTGCAGGACGGCGAGTTCATCTACAAG<br>GTGAAGCTGCGCGGCACCAACTTCCCCTCCGACGGCCCCGTAATGCAGAAGAAGACCATGGGC<br>TGGGAGGCCTCCTCCGAGCGGATGTACCCCGAGGACGGCGCCCTGAAGGGCGAGATCAAGATG<br>AGGCTGAAGCTGAAGGACGGCGGCCACTACGACGCTGAGGTCAAGACCACCTACAAGGCCAAG<br>AAGCCCGTGCAGCTGCCCGGCGCCTACATCGTCGGCATCAAGTTGGACATCACCTCCCACAACG<br>AGGACTACACCATCGTGGAACTGTACGAACGCGCCGAGGGCCGCCACTCCACCGGCGGCATGG<br>ACGAGCTGTACAAG**TAA** |
| mApple | AGATCTGTGAGCAAGGGCGAGGAGAATAACATGGCCATCATCAAGGAGTTCATGCGCTTCAAGGT<br>GCACATGGAGGGCTCCGTGAACGGCCACGAGTTCGAGATCGAGGGCGAGGGCGAGGGCCGCC<br>CCTACGAGGCCTTTCAGACCGCTAAGCTGAAGGTGACCAAGGGTGGCCCCCTGCCCTTCGCCT<br>GGGACATCCTGTCCCCTCAGTTCATGTACGGCTCCAAGGTCTACATTAAGCACCCAGCCGACATC<br>CCCGACTACTTCAAGCTGTCCTTCCCCGAGGGCTTCAGGTGGGAGCGCGTGATGAACTTCGAG<br>GACGGCGGCATTATTCACGTTAACCAGGACTCCTCCCTGCAGGACGGCGTGTTCATCTACAAGG<br>TGAAGCTGCGCGGCACCAACTTCCCCTCCGACGGCCCCGTAATGCAGAAGAAGACCATGGGCT<br>GGGAGGCCTCCGAGGAGCGGATGTACCCCGAGGACGGCGCCCTGAAGAGCGAGATCAAGAAG<br>AGGCTGAAGCTGAAGGACGGCGGCCACTACGCCGCCGAGGTCAAGACAGCCCGTGCAGCTAG<br>CCCGTGCAGCTGCCCGGCGCCTACATCGTCGACATCAAGTTGGACATCGTGTCCCACAACGAGG<br>ACTACACCATCGTGGAACAGTACGAACGCGCCGAGGGCCGCCACTCCACCGGCGGCATGGACG<br>AGCTGTACAAG**TAA** |
| mRuby2 | AGATCTGTGAGCAAGGGCGAAGAGCTGATCAAGGAAAATATGCGTATGAAGGTGGTCATGGAAG<br>GTTCGGTCAACGGCCACCAATTCAAATGCACAGGTGAAGGAGAAGGCAATCCGTACATGGGAAC<br>TCAAACCATGAGGATCAAAGTCATCGAGGGAGGACCCCTGCCATTTGCCTTTGACATTCTTGCCA<br>CGTCGTTCATGTATGGCAGCCGTACTTTTATCAAGTACCCGAAAGGCATTCCTGATTTCTTTAAACA<br>GTCCTTTCCTGAGGGTTTTACTTGGGAAAGAGTTACGAGATACGAAGATGGTGGAGTCGTCACCG |

| | |
|---|---|
| | TCATGCAGGACACCAGCCTTGAGGATGGCTGTCTCGTTTACCACGTCCAAGTCAGAGGGGTAAA<br>CTTTCCCTCCAATGGTCCCGTGATGCAGAAGAAGACCAAGGGTTGGGAGCCTAATACAGAGATG<br>ATGTATCCAGCAGATGGTGGTCTGAGGGGATACACTCATATGGCACTGAAAGTTGATGGTGGTGG<br>CCATCTGTCTTGCTCTTTCGTAACAACTTACAGGTCAAAAAAGACCGTCGGGAACATCAAGATGC<br>CCGGTATCCATGCCGTTGATCACCGCCTGGAAAGGTTAGAGGAAAGTGACAATGAAATGTTCGTA<br>GTACAACGCGAACACGCAGTTGCCAAGTTCGCCGGGCTTGGTGGTGGGATGGACGAGCTGTAC<br>AAG**TAA** |
| mScarlet-I | AGATCTGTGAGCAAGGGCGAGGCAGTGATCAAGGAGTTCATGCGGTTCAAGGTGCACATGGAG<br>GGCTCCATGAACGGCCACGAGTTCGAGATCGAGGGCGAGGGCGAGGGCCGCCCCTACGAGGG<br>CACCCAGACCGCCAAGCTGAAGGTGACCAAGGGTGGCCCCCTGCCCTTCTCCTGGGACATCCT<br>GTCCCCTCAGTTCATGTACGGCTCCAGGGCCTTCATCAAGCACCCCGCCGACATCCCCGACTAC<br>TATAAGCAGTCCTTCCCCGAGGGCTTCAAGTGGGAGCGCGTGATGAACTTCGAGGACGGCGGC<br>GCCGTGACCGTGACCCAGGACACCTCCCTGGAGGACGGCACCCTGATCTACAAGGTGAAGCTC<br>CGCGGCACCAACTTCCCTCCTGACGGCCCCGTAATGCAGAAGAAGACAATGGGCTGGGAAGCG<br>TCCACCGAGCGGTTGTACCCCGAGGACGGCGTGCTGAAGGGCGACATTAAGATGGCCCTGCGC<br>CTGAAGGACGGCGGCCGCTACCTGGCGGACTTCAAGACCACCTACAAGGCCAAGAAGCCCGTG<br>CAGATGCCCGGCGCCTACAACGTCGACCGCAAGTTGGACATCACCTCCCACAACGAGGACTACA<br>CCGTGGTGGAACAGTACGAACGCTCCGAGGGCCGCCACTCCACCGGCGGCATGGACGAGCTG<br>TACAAG**TAA** |
| mCherry | AGATCTGTGTCCAAGGGCGAGGAGGATAACATGGCCATCATCAAGGAGTTCATGCGCTTCAAGGT<br>GCACATGGAGGGCAGCGTGAACGGCCACGAGTTCGAGATTGAGGGCGAGGGCGAGGGACGCC<br>CATATGAGGGAACCCAGACCGCCAAGCTGAAAGTGACCAAGGGCGGACCCCTGCCCTTCGCCT<br>GGGATATTCTGAGCCCCCAGTTTATGTACGGCAGCAAGGCCTACGTGAAGCACCCCGCCGATATC<br>CCCGATTACCTGAAGCTGAGCTTCCCCGAGGGCTTCAAGTGGGAGCGCGTGATGAACTTCGAG<br>GATGGCGGCGTCGTGACCGTGACCCAGGATAGCAGTCTGCAGGATGGCGAGTTCATCTACAAAG<br>TGAAGCTGCGCGGCACCAACTTCCCCAGCGACGGACCAGTGATGCAGAAGAAGACCATGGGCT<br>GGGAGGCCAGCAGCGAGCGCATGTATCCAGAGGACGGCGCCCTGAAGGGCGAGATCAAGCAG<br>CGCCTGAAGCTGAAGGATGGCGGCCACTACGATGCCGAAGTGAAGACCACCTACAAGGCCAAG<br>AAGCCCGTGCAGCTGCCAGGCGCCTACAACGTGAACATCAAGCTGGATATCACCAGCCACAACG<br>AGGACTACACCATCGTGGAGCAGTACGAGCGCGCCGAGGGCCGCCATAGTACCGGCGGAATGG<br>ATGAGCTGTACAAG**TAG** |
| FusionRed-M | AGATCTGTGAGCGAGCTGATTAAGGAGAACATGCCCATGAAGCTGTACATGGAGGGCACCGTGA<br>ACAACCACCACTTCAAGTGCACATCCGAGGGCGAAGGCAAGCCCTACGAGGGCACCCAGACCA<br>TGAGAATCAAGGTCGTCGAGGGCGGCCCTCTCCCCTTCGCCTTCGACATCCTGGCTACCAGCTT<br>CATGTACGGCAGCAGAACCTTCATCAAGCACCCTCCGGGCATCCCCGACTTCTTTAAGCAGTCCT<br>TCCCTGAGGGCTTCACATGGGAGAGAGTCACCACATACGAAGACGGGGGCGTGCTGACCGCTA<br>CCCAGGACACCAGCCTCCAGGACGGCTGCCTCATCTACAACGTCAAGGTTAGAGGGGTGAACTT<br>CCCAGCCAACGGCCCTGTGATGCAGAAGAAAACACTCGGCTGGGAGGCCTCCACCGAGACGAT<br>GTACCCCGCTGACGGCGGCCTGGAAGGCGCATGTGACATGGCCCTGAAGCTCGTGGGCGGGG<br>GCCACATGATCTGCAACCTTGAGACCACATACAGATCCAAGAAACCCGCTACGAACCTCAAGATG<br>CCCGGCGTCTACAACGTGGACCACAGACTGGAAAGAATCAAGGAGGCCGACGATGAGACCTAC<br>GTCGAGCAGCACGAGGTGGCTGTGGCCAGATACTCTACTGGTGGCGCTGGTGATGGAGGTAAA**T**<br>**GA** |

**Supplementary Table 2:** Sequences for the selected red fluorescent proteins.

### Supplementary Figures

A

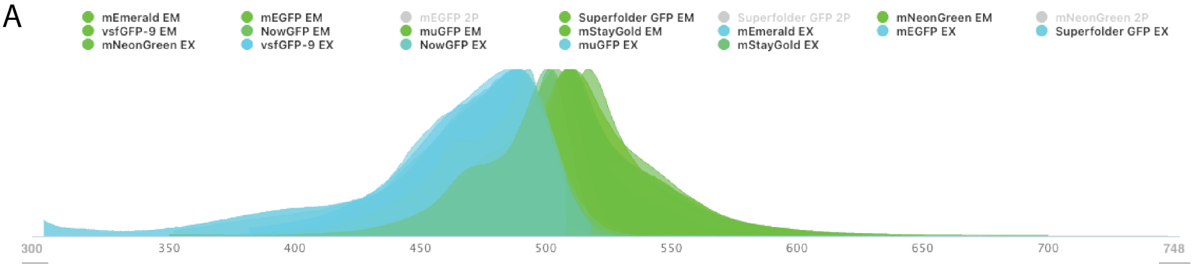


Attribute Comparison

| Name | $\lambda_{ex}$ | $\lambda_{em}$ | Stokes | EC | QY | Brightness | pKa | Aggregation | Maturation | Lifetime | kDa |
|---|---|---|---|---|---|---|---|---|---|---|---|
| mEmerald | 487 | 509 | 22 | 57,500 | 0.68 | 39.1 | 6.0 | m | 11.2 | | 26.95 |
| mEGFP | 488 | 507 | 19 | 56,000 | 0.6 | 33.6 | 6.0 | m | | | 27.00 |
| Superfolder GFP | 488 | 510 | 22 | 83,300 | 0.65 | 54.15 | | wd | 13.6 | 3.1 | 26.78 |
| SGFP2 | 495 | 512 | 17 | 46,000 | 0.7 | 32.2 | 5.9 | m | | | 26.89 |
| mNeonGreen | 506 | 517 | 11 | 116,000 | 0.8 | 92.8 | 5.7 | m | 10.0 | 3.1 | 26.65 |
| vsfGFP-9 | 485 | 510 | 25 | 104,958 | 0.7 | 73.47 | 4.75 | m | | | 39.46 |
| NowGFP | 494 | 502 | 8 | 56,700 | 0.76 | 43.09 | 6.2 | m | | 5.1 | 26.69 |
| muGFP | 490 | 508 | 18 | 117,000 | 0.45 | 52.65 | | m | | | 26.78 |
| mStayGold | 499 | 510 | 11 | 164,000 | 0.83 | 136.12 | 4.8 | m | | 2.79 | 25.58 |

B

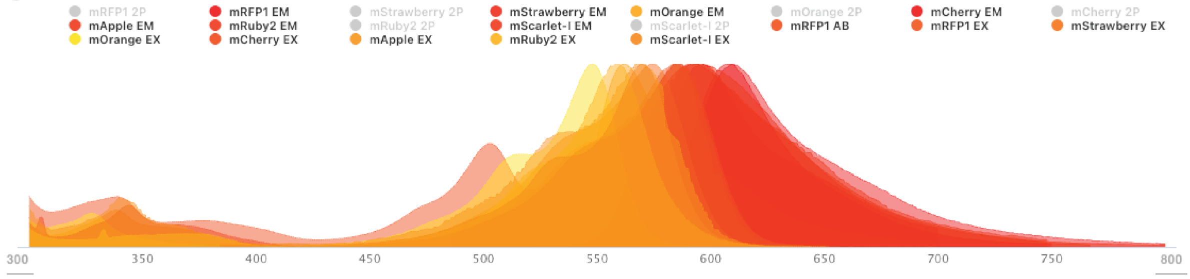


Attribute Comparison

| Name | $\lambda_{ex}$ | $\lambda_{em}$ | Stokes | EC | QY | Brightness | pKa | Aggregation | Maturation | Lifetime | kDa |
|---|---|---|---|---|---|---|---|---|---|---|---|
| mRFP1 | 584 | 607 | 23 | 50,000 | 0.25 | 12.5 | 4.5 | m | 60.0 | | 25.42 |
| mStrawberry | 574 | 596 | 22 | 90,000 | 0.29 | 26.1 | 4.5 | m | 50.0 | | 26.64 |
| mOrange | 548 | 562 | 14 | 71,000 | 0.69 | 48.99 | 6.5 | m | 150.0 | | 26.74 |
| mCherry | 587 | 610 | 23 | 72,000 | 0.22 | 15.84 | 4.5 | m | 15.0 | 1.4 | 26.72 |
| mApple | 568 | 592 | 24 | 75,000 | 0.49 | 36.75 | 6.5 | m | 30.0 | 2.9 | 26.96 |
| mRuby2 | 559 | 600 | 41 | 113,000 | 0.38 | 42.94 | 5.3 | m | 150.0 | 2.5 | 26.49 |
| mScarlet-I | 569 | 593 | 24 | 104,000 | 0.54 | 56.16 | 5.4 | m | 36.0 | 3.1 | 26.36 |
| FusionRed-M | 571 | 594 | 23 | 71,100 | 0.34 | 24.17 | | m | | 2.1 | 25.60 |

**Figure S1**: Fluorescent protein characterisation from FP Base[1]. Comparison of (**A**) green and (**B**) red fluorescent proteins. $\lambda_{ex}$ and $\lambda_{em}$ refer to excitation and emission wavelengths. Stokes = Stokes Shift. EC = Extinction Coefficient. QY = Quantum Yield. Brightness = intensity per fluorophore emission = EC x QY. pKa = pH at which half of the fluorophore is deprotonated. Aggregation, m = monomer, wd = weak dimer. Maturation = estimated folding time (mins), defined by time to reach half-maximal value. Lifetime = inverse of de-excitation rate (ns), and describes the time for the fluorophore to relax to the ground state. kDa = Molecular size.

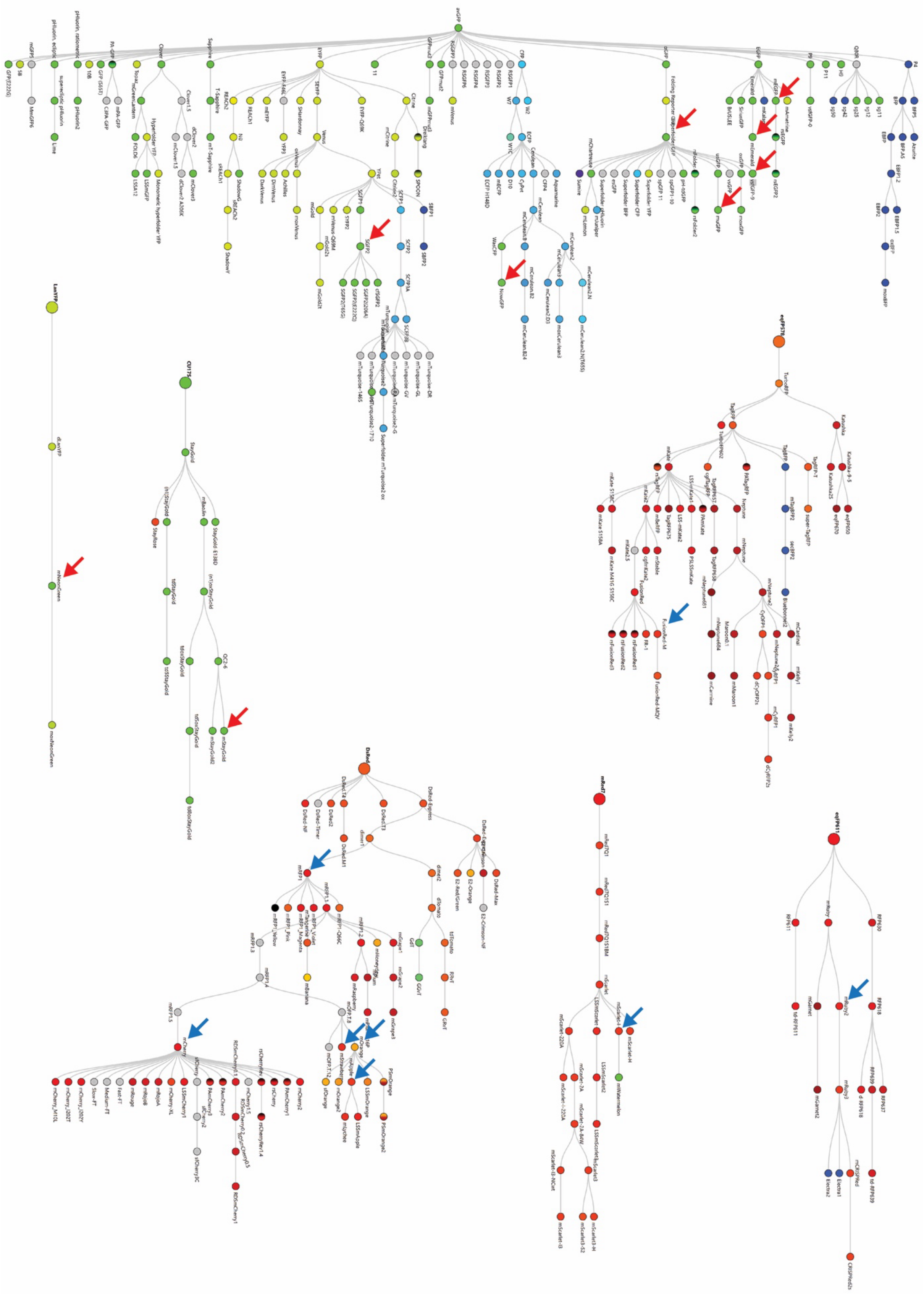

**Figure S2**: Fluorophore evolution trees for both red and green fluorophores. Arrows highlight selected fluorophores in this study. Figure generated using https://www.fpbase.org/lineage/ [1].

A

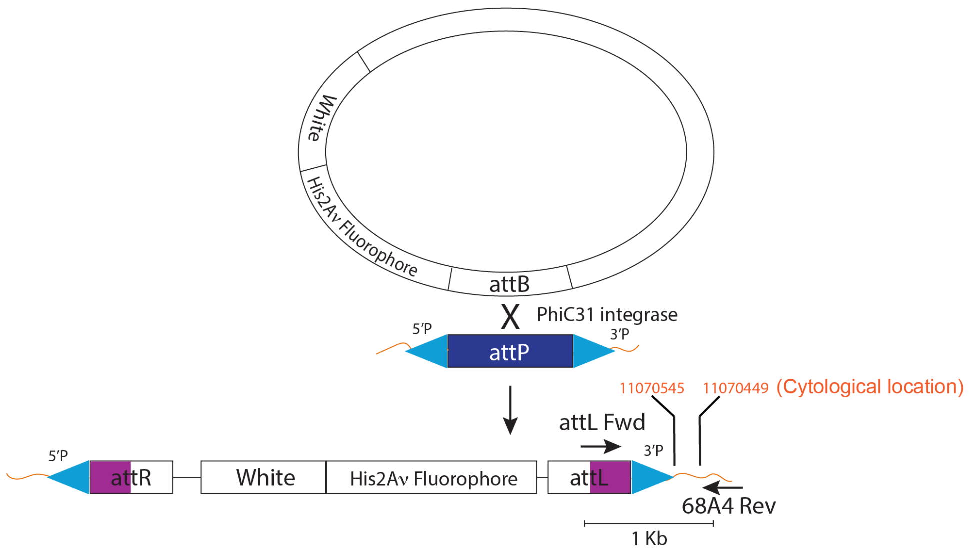


B

```
                                                part of attL       P element 3' end

mEmerald        --AAATGGGGGTGGGTCGCCGAACATGACACAAGGGGTTGTGACCGGGGTGGACACGTAC      58
mEGFP           --AAAAGGGGGTGGGTCGCCGAACATGACACAAGGGGTTGTGACCGGGGTGGACACGTAC      58
sfGFP           ------ATGGTCGCCGAAAATGAACACAAAGGGGGTTTGTGACCGGGGTGGGACACGTAC      54
sGFP2           ----AAGGGGGTGGGTCGCCGAACATGACACAAGGGGTTGTGACCGGGGTGGACACGTAC      56
mNeonGreen      AAATGGGGGTGGGGTCGCCGAAATGACCACAAGGGGGTTGTGACCGGGGTGGACACGTAC      60
vsfGFP-9        ---------------------------AAGGGGGTTGTGGACCGGGGGTGGGACACGTAC      33
NowGFP          ---NANNNGGGGNGGTCGCCAGAACTGACACAAGGGGTTGTGACCGGGGTGGACACGTAC      57
muGFP           --AAAATGGCT--AGGTCGCCGAACTGACACAAGGGGTTGTGACCGGGGTGGACACGTAC      56
mRFP1           ----ATGGGGTGGGTCGCCGAACATGAACACAAGGGGTTGTGACCGGGGTGGACACGTAC      56
mOrange         -----------TAGGGCGCCGACATGACACAAGGGGGTTGTGACCGGGGTGGACACGTAC      49
mStrawberry     -----ATGGGGAGGGTCGCCGAACATGACACAAGGGGTTGTGACCGGGGTGGACACGTAC      55
mApple          ----AGGGGGTGGGTCGCCGAACATGAACACAAGGGGTTGTGACCGGGGTGGACACGTAC      56
mRuby2          ---AATGGGGTGGGTCGCCGAACATGAACACAAGGGGTTGTGACCGGGGTGGACACGTAC      57
mScarlet-I      ------------TAGGGCGCCGACTGAACACAAGGGGTTGTGACCGGGGTGGACACGTAC      48
mCherry         -----------TAGGGCGCCGACATGACACAAGGGGGTTGTGACCGGGGTGGACACGTAC      49
FusionRedM      ---NAGGGGGTAGGGTCGCCGAACTGAACACAAGGGGTTGTGACCGGGGTGGACACGTAC      57

                                   P element 3' end

mEmerald        GCGGGTGCTTACGACCGTCAGTCGCGCGAGCGCGACTAGGGGCGAGCTCGAATTGGCGGC      118
mEGFP           GCGGGTGCTTACGACCGTCAGTCGCGCGAGCGCGACTAGGGGCGAGCTCGAATTGGCGGC      118
sfGFP           GCGGGTGCTTACGACCGTCAGTCGCGCGAGCGCGACTAGGGGCGAGCTCGAATTGGCGGC      114
sGFP2           GCGGGTGCTTACGACCGTCAGTCGCGCGAGCGCGACTAGGGGCGAGCTCGAATTGGCGGC      116
mNeonGreen      GCGGGTGCTTACGACCGTCAGTCGCGCGAGCGCGACTAGGGGCGAGCTCGAATTGGCGGC      120
vsfGFP-9        GCGGGTGCTTACGACCGTCAGTCGCGCGAGCGCGACTAGGGGCGAGCTCGAATTGGCGGC      93
NowGFP          GCGGGTGCTTACGACCGTCAGTCGCGCGAGCGCGACTAGGGGCGAGCTCGAATTGGCGGC      117
muGFP           GCGGGTGCTTACGACCGTCAGTCGCGCGAGCGCGACTAGGGGCGAGCTCGAATTGGCGGC      116
mRFP1           GCGGGTGCTTACGACCGTCAGTCGCGCGAGCGCGACTAGGGGCGAGCTCGAATTGGCGGC      116
mOrange         GCGGGTGCTTACGACCGTCAGTCGCGCGAGCGCGACTAGGGGCGAGCTCGAATTGGCGGC      109
mStrawberry     GCGGGTGCTTACGACCGTCAGTCGCGCGAGCGCGACTAGGGGCGAGCTCGAATTGGCGGC      115
mApple          GCGGGTGCTTACGACCGTCAGTCGCGCGAGCGCGACTAGGGGCGAGCTCGAATTGGCGGC      116
mRuby2          GCGGGTGCTTACGACCGTCAGTCGCGCGAGCGCGACTAGGGGCGAGCTCGAATTGGCGGC      117
mScarlet-I      GCGGGTGCTTACGACCGTCAGTCGCGCGAGCGCGACTAGGGGCGAGCTCGAATTGGCGGC      108
mCherry         GCGGGTGCTTACGACCGTCAGTCGCGCGAGCGCGACTAGGGGCGAGCTCGAATTGGCGGC      109
FusionRedM      GCGGGTGCTTACGACCGTCAGTCGCGCGAGCGCGACTAGGGGCGAGCTCGAATTGGCGGC      117
```

P element 3’ end

| | | |
|---|---|---|
| mEmerald | CGCCGGCGGCCGCCAATTCGGCTGCTGCTCTAAACGACGCATTTCGTACTCCAAAGTACG | 178 |
| mEGFP | CGCCGGCGGCCGCCAATTCGGCTGCTGCTCTAAACGACGCATTTCGTACTCCAAAGTACG | 178 |
| sfGFP | CGCCGGCGGCCGCCAATTCGGCTGCTGCTCTAAACGACGCATTTCGTACTCCAAAGTACG | 174 |
| sGFP2 | CGCCGGCGGCCGCCAATTCGGCTGCTGCTCTAAACGACGCATTTCGTACTCCAAAGTACG | 176 |
| mNeonGreen | CGCCGGCGGCCGCCAATTCGGCTGCTGCTCTAAACGACGCATTTCGTACTCCAAAGTACG | 180 |
| vsfGFP-9 | CGCCGGCGGCCGCCAATTCGGCTGCTGCTCTAAACGACGCATTTCGTACTCCAAAGTACG | 153 |
| NowGFP | CGCCGGCGGCCGCCAATTCGGCTGCTGCTCTAAACGACGCATTTCGTACTCCAAAGTACG | 177 |
| muGFP | CGCCGGCGGCCGCCAATTCGGCTGCTGCTCTAAACGACGCATTTCGTACTCCAAAGTACG | 176 |
| mRFP1 | CGCCGGCGGCCGCCAATTCGGCTGCTGCTCTAAACGACGCATTTCGTACTCCAAAGTACG | 176 |
| mOrange | CGCCGGCGGCCGCCAATTCGGCTGCTGCTCTAAACGACGCATTTCGTACTCCAAAGTACG | 169 |
| mStrawberry | CGCCGGCGGCCGCCAATTCGGCTGCTGCTCTAAACGACGCATTTCGTACTCCAAAGTACG | 175 |
| mApple | CGCCGGCGGCCGCCAATTCGGCTGCTGCTCTAAACGACGCATTTCGTACTCCAAAGTACG | 176 |
| mRuby2 | CGCCGGCGGCCGCCAATTCGGCTGCTGCTCTAAACGACGCATTTCGTACTCCAAAGTACG | 177 |
| mScarlet-I | CGCCGGCGGCCGCCAATTCGGCTGCTGCTCTAAACGACGCATTTCGTACTCCAAAGTACG | 168 |
| mCherry | CGCCGGCGGCCGCCAATTCGGCTGCTGCTCTAAACGACGCATTTCGTACTCCAAAGTACG | 169 |
| FusionRedM | CGCCGGCGGCCGCCAATTCGGCTGCTGCTCTAAACGACGCATTTCGTACTCCAAAGTACG | 177 |

P element 3’ end

| | | |
|---|---|---|
| mEmerald | AATTTTTTCCCTCAAGCTCTTATTTTCATTAAACAATGAACAGGACCTAACGCACAGTCA | 238 |
| mEGFP | AATTTTTTCCCTCAAGCTCTTATTTTCATTAAACAATGAACAGGACCTAACGCACAGTCA | 238 |
| sfGFP | AATTTTTTCCCTCAAGCTCTTATTTTCATTAAACAATGAACAGGACCTAACGCACAGTCA | 234 |
| sGFP2 | AATTTTTTCCCTCAAGCTCTTATTTTCATTAAACAATGAACAGGACCTAACGCACAGTCA | 236 |
| mNeonGreen | AATTTTTTCCCTCAAGCTCTTATTTTCATTAAACAATGAACAGGACCTAACGCACAGTCA | 240 |
| vsfGFP-9 | AATTTTTTCCCTCAAGCTCTTATTTTCATTAAACAATGAACAGGACCTAACGCACAGTCA | 213 |
| NowGFP | AATTTTTTCCCTCAAGCTCTTATTTTCATTAAACAATGAACAGGACCTAACGCACAGTCA | 237 |
| muGFP | AATTTTTTCCCTCAAGCTCTTATTTTCATTAAACAATGAACAGGACCTAACGCACAGTCA | 236 |
| mRFP1 | AATTTTTTCCCTCAAGCTCTTATTTTCATTAAACAATGAACAGGACCTAACGCACAGTCA | 236 |
| mOrange | AATTTTTTCCCTCAAGCTCTTATTTTCATTAAACAATGAACAGGACCTAACGCACAGTCA | 229 |
| mStrawberry | AATTTTTTCCCTCAAGCTCTTATTTTCATTAAACAATGAACAGGACCTAACGCACAGTCA | 235 |
| mApple | AATTTTTTCCCTCAAGCTCTTATTTTCATTAAACAATGAACAGGACCTAACGCACAGTCA | 236 |
| mRuby2 | AATTTTTTCCCTCAAGCTCTTATTTTCATTAAACAATGAACAGGACCTAACGCACAGTCA | 237 |
| mScarlet-I | AATTTTTTCCCTCAAGCTCTTATTTTCATTAAACAATGAACAGGACCTAACGCACAGTCA | 228 |
| mCherry | AATTTTTTCCCTCAAGCTCTTATTTTCATTAAACAATGAACAGGACCTAACGCACAGTCA | 229 |
| FusionRedM | AATTTTTTCCCTCAAGCTCTTATTTTCATTAAACAATGAACAGGACCTAACGCACAGTCA | 237 |

P element 3’ end

| | | |
|---|---|---|
| mEmerald | CGTTATTGTTTACATAAATGATTTTTTTTACTATTCAAACTTACTCTGTTTGTGTACTCC | 298 |
| mEGFP | CGTTATTGTTTACATAAATGATTTTTTTTACTATTCAAACTTACTCTGTTTGTGTACTCC | 298 |
| sfGFP | CGTTATTGTTTACATAAATGATTTTTTTTACTATTCAAACTTACTCTGTTTGTGTACTCC | 294 |
| sGFP2 | CGTTATTGTTTACATAAATGATTTTTTTTACTATTCAAACTTACTCTGTTTGTGTACTCC | 296 |
| mNeonGreen | CGTTATTGTTTACATAAATGATTTTTTTTACTATTCAAACTTACTCTGTTTGTGTACTCC | 300 |
| vsfGFP-9 | CGTTATTGTTTACATAAATGATTTTTTTTACTATTCAAACTTACTCTGTTTGTGTACTCC | 273 |
| NowGFP | CGTTATTGTTTACATAAATGATTTTTTTTAATATTCAAACTTACTCTGTTTGTGTACTCC | 297 |
| muGFP | CGTTATTGTTTACATAAATGATTTTTTTTACTATTCAAACTTACTCTGTTTGTGTACTCC | 296 |
| mRFP1 | CGTTATTGTTTACATAAATGATTTTTTTTACTATTCAAACTTACTCTGTTTGTGTACTCC | 296 |
| mOrange | CGTTATTGTTTACATAAATGATTTTTTTTAATATTCAAACTTACTCTGTTTGTGTACTCC | 289 |
| mStrawberry | CGTTATTGTTTACATAAATGATTTTTTTTAATATTCAAACTTACTCTGTTTGTGTACTCC | 295 |
| mApple | CGTTATTGTTTACATAAATGATTTTTTTTAATATTCAAACTTACTCTGTTTGTGTACTCC | 296 |
| mRuby2 | CGTTATTGTTTACATAAATGATTTTTTTTAATATTCAAACTTACTCTGTTTGTGTACTCC | 297 |
| mScarlet-I | CGTTATTGTTTACATAAATGATTTTTTTTACTATTCAAACTTACTCTGTTTGTGTACTCC | 288 |
| mCherry | CGTTATTGTTTACATAAATGATTTTTTTTACTATTCAAACTTACTCTGTTTGTGTACTCC | 289 |
| FusionRedM | CGTTATTGTTTACATAAATGATTTTTTTTACTATTCAAACTTACTCTGTTTGTGTACTCC | 297 |

P element 3’ end

| | | |
|---|---|---|
| mEmerald | CACTGGTATAGCCTTCTTTTATCTTTTCTGGTTCAGGCTCTATCACTTTACTAGGTACGG | 358 |
| mEGFP | CACTGGTATAGCCTTCTTTTATCTTTTCTGGTTCAGGCTCTATCACTTTACTAGGTACGG | 358 |
| sfGFP | CACTGGTATAGCCTTCTTTTATCTTTTCTGGTTCAGGCTCTATCACTTTACTAGGTACGG | 354 |
| sGFP2 | CACTGGTATAGCCTTCTTTTATCTTTTCTGGTTCAGGCTCTATCACTTTACTAGGTACGG | 356 |
| mNeonGreen | CACTGGTATAGCCTTCTTTTATCTTTTCTGGTTCAGGCTCTATCACTTTACTAGGTACGG | 360 |
| vsfGFP-9 | CACTGGTATAGCCTTCTTTTATCTTTTCTGGTTCAGGCTCTATCACTTTACTAGGTACGG | 333 |
| NowGFP | CACTGGTATAGCCTTCTTTTATCTTTTCTGGTTCAGGCTCTATCACTTTACTAGGTACGG | 357 |
| muGFP | CACTGGTATAGCCTTCTTTTATCTTTTCTGGTTCAGGCTCTATCACTTTACTAGGTACGG | 356 |
| mRFP1 | CACTGGTATAGCCTTCTTTTATCTTTTCTGGTTCAGGCTCTATCACTTTACTAGGTACGG | 356 |
| mOrange | CACTGGTATAGCCTTCTTTTATCTTTTCTGGTTCAGGCTCTATCACTTTACTAGGTACGG | 349 |
| mStrawberry | CACTGGTATAGCCTTCTTTTATCTTTTCTGGTTCAGGCTCTATCACTTTACTAGGTACGG | 355 |
| mApple | CACTGGTATAGCCTTCTTTTATCTTTTCTGGTTCAGGCTCTATCACTTTACTAGGTACGG | 356 |
| mRuby2 | CACTGGTATAGCCTTCTTTTATCTTTTCTGGTTCAGGCTCTATCACTTTACTAGGTACGG | 357 |
| mScarlet-I | CACTGGTATAGCCTTCTTTTATCTTTTCTGGTTCAGGCTCTATCACTTTACTAGGTACGG | 348 |
| mCherry | CACTGGTATAGCCTTCTTTTATCTTTTCTGGTTCAGGCTCTATCACTTTACTAGGTACGG | 349 |
| FusionRedM | CACTGGTATAGCCTTCTTTTATCTTTTCTGGTTCAGGCTCTATCACTTTACTAGGTACGG | 357 |

P element 3' end

```
mEmerald        CATCTGCGTTGAGTCGCCTCCTTTTAAATGTCTGACCTTTTGCAGGTGCAGCCTTCCACT   418
mEGFP           CATCTGCGTTGAGTCGCCTCCTTTTAAATGTCTGACCTTTTGCAGGTGCAGCCTTCCACT   418
sfGFP           CATCTGCGTTGAGTCGCCTCCTTTTAAATGTCTGACCTTTTGCAGGTGCAGCCTTCCACT   414
sGFP2           CATCTGCGTTGAGTCGCCTCCTTTTAAATGTCTGACCTTTTGCAGGTGCAGCCTTCCACT   416
mNeonGreen      CATCTGCGTTGAGTCGCCTCCTTTTAAATGTCTGACCTTTTGCAGGTGCAGCCTTCCACT   420
vsfGFP-9        CATCTGCGTTGAGTCGCCTCCTTTTAAATGTCTGACCTTTTGCAGGTGCAGCCTTCCACT   393
NowGFP          CATCTGCGTTGAGTCGCCTCCTTTTAAATGTCTGACCTTTTGCAGGTGCAGCCTTCCACT   417
muGFP           CATCTGCGTTGAGTCGCCTCCTTTTAAATGTCTGACCTTTTGCAGGTGCAGCCTTCCACT   416
mRFP1           CATCTGCGTTGAGTCGCCTCCTTTTAAATGTCTGACCTTTTGCAGGTGCAGCCTTCCACT   416
mOrange         CATCTGCGTTGAGTCGCCTCCTTTTAAATGTCTGACCTTTTGCAGGTGCAGCCTTCCACT   409
mStrawberry     CATCTGCGTTGAGTCGCCTCCTTTTAAATGTCTGACCTTTTGCAGGTGCAGCCTTCCACT   415
mApple          CATCTGCGTTGAGTCGCCTCCTTTTAAATGTCTGACCTTTTGCAGGTGCAGCCTTCCACT   416
mRuby2          CATCTGCGTTGAGTCGCCTCCTTTTAAATGTCTGACCTTTTGCAGGTGCAGCCTTCCACT   417
mScarlet-I      CATCTGCGTTGAGTCGCCTCCTTTTAAATGTCTGACCTTTTGCAGGTGCAGCCTTCCACT   408
mCherry         CATCTGCGTTGAGTCGCCTCCTTTTAAATGTCTGACCTTTTGCAGGTGCAGCCTTCCACT   409
FusionRedM      CATCTGCGTTGAGTCGCCTCCTTTTAAATGTCTGACCTTTTGCAGGTGCAGCCTTCCACT   417
```

P element 3' end

```
mEmerald        GCGAATCATTAAAGTGGGTATCACAAATTTGGGAGTTTTCACCAAGGCTGCACCCAAGGC   478
mEGFP           GCGAATCATTAAAGTGGGTATCACAAATTTGGGAGTTTTCACCAAGGCTGCACCCAAGGC   478
sfGFP           GCGAATCATTAAAGTGGGTATCACAAATTTGGGAGTTTTCACCAAGGCTGCACCCAAGGC   474
sGFP2           GCGAATCATTAAAGTGGGTATCACAAATTTGGGAGTTTTCACCAAGGCTGCACCCAAGGC   476
mNeonGreen      GCGAATCATTAAAGTGGGTATCACAAATTTGGGAGTTTTCACCAAGGCTGCACCCAAGGC   480
vsfGFP-9        GCGAATCATTAAAGTGGGTATCACAAATTTGGGAGTTTTCACCAAGGCTGCACCCAAGGC   453
NowGFP          GCGAATCATTAAAGTGGGTATCACAAATTTGGGAGTTTTCACCAAGGCTGCACCCAAGGC   477
muGFP           GCGAATCATTAAAGTGGGTATCACAAATTTGGGAGTTTTCACCAAGGCTGCACCCAAGGC   476
mRFP1           GCGAATCATTAAAGTGGGTATCACAAATTTGGGAGTTTTCACCAAGGCTGCACCCAAGGC   476
mOrange         GCGAATCATTAAAGTGGGTATCACAAATTTGGGAGTTTTCACCAAGGCTGCACCCAAGGC   469
mStrawberry     GCGAATCATTAAAGTGGGTATCACAAATTTGGGAGTTTTCACCAAGGCTGCACCCAAGGC   475
mApple          GCGAATCATTAAAGTGGGTATCACAAATTTGGGAGTTTTCACCAAGGCTGCACCCAAGGC   476
mRuby2          GCGAATCATTAAAGTGGGTATCACAAATTTGGGAGTTTTCACCAAGGCTGCACCCAAGGC   477
mScarlet-I      GCGAATCATTAAAGTGGGTATCACAAATTTGGGAGTTTTCACCAAGGCTGCACCCAAGGC   468
mCherry         GCGAATCATTAAAGTGGGTATCACAAATTTGGGAGTTTTCACCAAGGCTGCACCCAAGGC   469
FusionRedM      GCGAATCATTAAAGTGGGTATCACAAATTTGGGAGTTTTCACCAAGGCTGCACCCAAGGC   477
```

P element 3' end

```
mEmerald        TCTGCTCCCACAATTTTCTCTTAATAGCACACTTCGGCACGTGAATTAATTTTACTCCAG   538
mEGFP           TCTGCTCCCACAATTTTCTCTTAATAGCACACTTCGGCACGTGAATTAATTTTACTCCAG   538
sfGFP           TCTGCTCCCACAATTTTCTCTTAATAGCACACTTCGGCACGTGAATTAATTTTACTCCAG   534
sGFP2           TCTGCTCCCACAATTTTCTCTTAATAGCACACTTCGGCACGTGAATTAATTTTACTCCAG   536
mNeonGreen      TCTGCTCCCACAATTTTCTCTTAATAGCACACTTCGGCACGTGAATTAATTTTACTCCAG   540
vsfGFP-9        TCTGCTCCCACAATTTTCTCTTAATAGCACACTTCGGCACGTGAATTAATTTTACTCCAG   513
NowGFP          TCTGCTCCCACAATTTTCTCTTAATAGCACACTTCGGCACGTGAATTAATTTTACTCCAG   537
muGFP           TCTGCTCCCACAATTTTCTCTTAATAGCACACTTCGGCACGTGAATTAATTTTACTCCAG   536
mRFP1           TCTGCTCCCACAATTTTCTCTTAATAGCACACTTCGGCACGTGAATTAATTTTACTCCAG   536
mOrange         TCTGCTCCCACAATTTTCTCTTAATAGCACACTTCGGCACGTGAATTAATTTTACTCCAG   529
mStrawberry     TCTGCTCCCACAATTTTCTCTTAATAGCACACTTCGGCACGTGAATTAATTTTACTCCAG   535
mApple          TCTGCTCCCACAATTTTCTCTTAATAGCACACTTCGGCACGTGAATTAATTTTACTCCAG   536
mRuby2          TCTGCTCCCACAATTTTCTCTTAATAGCACACTTCGGCACGTGAATTAATTTTACTCCAG   537
mScarlet-I      TCTGCTCCCACAATTTTCTCTTAATAGCACACTTCGGCACGTGAATTAATTTTACTCCAG   528
mCherry         TCTGCTCCCACAATTTTCTCTTAATAGCACACTTCGGCACGTGAATTAATTTTACTCCAG   529
FusionRedM      TCTGCTCCCACAATTTTCTCTTAATAGCACACTTCGGCACGTGAATTAATTTTACTCCAG   537
```

P element 3' end

```
mEmerald        TCACAGCTTTGCAGCAAAATTTGCAATATTTCATTTTTTTTTATTCCACGTAAGGGTTAA   598
mEGFP           TCACAGCTTTGCAGCAAAATTTGCAATATTTCATTTTTTTTTATTCCACGTAAGGGTTAA   598
sfGFP           TCACAGCTTTGCAGCAAAATTTGCAATATTTCATTTTTTTTTATTCCACGTAAGGGTTAA   594
sGFP2           TCACAGCTTTGCAGCAAAATTTGCAATATTTCATTTTTTTTTATTCCACGTAAGGGTTAA   596
mNeonGreen      TCACAGCTTTGCAGCAAAATTTGCAATATTTCATTTTTTTTTATTCCACGTAAGGGTTAA   600
vsfGFP-9        TCACAGCTTTGCAGCAAAATTTGCAATATTTCATTTTTTTTTATTCCACGTAAGGGTTAA   573
NowGFP          TCACAGCTTTGCAGCAAAATTTGCAATATTTCATTTTTTTTTATTCCACGTAAGGGTTAA   597
muGFP           TCACAGCTTTGCAGCAAAATTTGCAATATTTCATTTTTTTTTATTCCACGTAAGGGTTAA   596
mRFP1           TCACAGCTTTGCAGCAAAATTTGCAATATTTCATTTTTTTTTATTCCACGTAAGGGTTAA   596
mOrange         TCACAGCTTTGCAGCAAAATTTGCAATATTTCATTTTTTTTTATTCCACGTAAGGGTTAA   589
mStrawberry     TCACAGCTTTGCAGCAAAATTTGCAATATTTCATTTTTTTTTATTCCACGTAAGGGTTAA   595
mApple          TCACAGCTTTGCAGCAAAATTTGCAATATTTCATTTTTTTTTATTCCACGTAAGGGTTAA   596
mRuby2          TCACAGCTTTGCAGCAAAATTTGCAATATTTCATTTTTTTTTATTCCACGTAAGGGTTAA   597
mScarlet-I      TCACAGCTTTGCAGCAAAATTTGCAATATTTCATTTTTTTTTATTCCACGTAAGGGTTAA   588
mCherry         TCACAGCTTTGCAGCAAAATTTGCAATATTTCATTTTTTTTTATTCCACGTAAGGGTTAA   589
FusionRedM      TCACAGCTTTGCAGCAAAATTTGCAATATTTCATTTTTTTTTATTCCACGTAAGGGTTAA   597
```

P element 3' end

```
mEmerald      TGTTTTCAAAAAAAAATTCGTCCGCACACAACCTTTCCTCTCAACAAGCAAACGTGCACT   658
mEGFP         TGTTTTCAAAAAAAAATTCGTCCGCACACAACCTTTCCTCTCAACAAGCAAACGTGCACT   658
sfGFP         TGTTTTCAAAAAAAAATTCGTCCGCACACAACCTTTCCTCTCAACAAGCAAACGTGCACT   654
sGFP2         TGTTTTCAAAAAAAAATTCGTCCGCACACAACCTTTCCTCTCAACAAGCAAACGTGCACT   656
mNeonGreen    TGTTTTCAAAAAAAAATTCGTCCGCACACAACCTTTCCTCTCAACAAGCAAACGTGCACT   660
vsfGFP-9      TGTTTTCAAAAAAAAATTCGTCCGCACACAACCTTTCCTCTCAACAAGCAAACGTGCACT   633
NowGFP        TGTTTTCAAAAAAAAATTCGTCCGCACACAACCTTTCCTCTCAACAAGCAAACGTGCACT   657
muGFP         TGTTTTCAAAAAAAAATTCGTCCGCACACAACCTTTCCTCTCAACAAGCAAACGTGCACT   656
mRFP1         TGTTTTCAAAAAAAAATTCGTCCGCACACAACCTTTCCTCTCAACAAGCAAACGTGCACT   656
mOrange       TGTTTTCAAAAAAAAATTCGTCCGCACACAACCTTTCCTCTCAACAAGCAAACGTGCACT   649
mStrawberry   TGTTTTCAAAAAAAAATTCGTCCGCACACAACCTTTCCTCTCAACAAGCAAACGTGCACT   655
mApple        TGTTTTCAAAAAAAAATTCGTCCGCACACAACCTTTCCTCTCAACAAGCAAACGTGCACT   656
mRuby2        TGTTTTCAAAAAAAAATTCGTCCGCACACAACCTTTCCTCTCAACAAGCAAACGTGCACT   657
mScarlet-I    TGTTTTCAAAAAAAAATTCGTCCGCACACAACCTTTCCTCTCAACAAGCAAACGTGCACT   648
mCherry       TGTTTTCAAAAAAAAATTCGTCCGCACACAACCTTTCCTCTCAACAAGCAAACGTGCACT   649
FusionRedM    TGTTTTCAAAAAAAAATTCGTCCGCACACAACCTTTCCTCTCAACAAGCAAACGTGCACT   657
```

P element 3' end

```
mEmerald      GAATTTAAGTGTATACTTCGGTAAGCTTCGGCTATCGACGGGACCACCTTATGTTATTTC   718
mEGFP         GAATTTAAGTGTATACTTCGGTAAGCTTCGGCTATCGACGGGACCACCTTATGTTATTTC   718
sfGFP         GAATTTAAGTGTATACTTCGGTAAGCTTCGGCTATCGACGGGACCACCTTATGTTATTTC   714
sGFP2         GAATTTAAGTGTATACTTCGGTAAGCTTCGGCTATCGACGGGACCACCTTATGTTATTTC   716
mNeonGreen    GAATTTAAGTGTATACTTCGGTAAGCTTCGGCTATCGACGGGACCACCTTATGTTATTTC   720
vsfGFP-9      GAATTTAAGTGTATACTTCGGTAAGCTTCGGCTATCGACGGGACCACCTTATGTTATTTC   693
NowGFP        GAATTTAAGTGTATACTTCGGTAAGCTTCGGCTATCGACGGGACCACCTTATGTTATTTC   717
muGFP         GAATTTAAGTGTATACTTCGGTAAGCTTCGGCTATCGACGGGACCACCTTATGTTATTTC   716
mRFP1         GAATTTAAGTGTATACTTCGGTAAGCTTCGGCTATCGACGGGACCACCTTATGTTATTTC   716
mOrange       GAATTTAAGTGTATACTTCGGTAAGCTTCGGCTATCGACGGGACCACCTTATGTTATTTC   709
mStrawberry   GAATTTAAGTGTATACTTCGGTAAGCTTCGGCTATCGACGGGACCACCTTATGTTATTTC   715
mApple        GAATTTAAGTGTATACTTCGGTAAGCTTCGGCTATCGACGGGACCACCTTATGTTATTTC   716
mRuby2        GAATTTAAGTGTATACTTCGGTAAGCTTCGGCTATCGACGGGACCACCTTATGTTATTTC   717
mScarlet-I    GAATTTAAGTGTATACTTCGGTAAGCTTCGGCTATCGACGGGACCACCTTATGTTATTTC   708
mCherry       GAATTTAAGTGTATACTTCGGTAAGCTTCGGCTATCGACGGGACCACCTTATGTTATTTC   709
FusionRedM    GAATTTAAGTGTATACTTCGGTAAGCTTCGGCTATCGACGGGACCACCTTATGTTATTTC   717
```

11070545 ▼ Endogenous region Cytological location 68A4

```
mEmerald      ATCATGATATGGACGAGCTGCCAGCGCAGCTTTCGCCAAAGCTTTTGTTCTAGTGCCAGT   778
mEGFP         ATCATGATATGGACGAGCTGCCAGCGCAGCTTTCGCCAAAGCTTTTGTTCTAGTGCCAGT   778
sfGFP         ATCATGATATGGACGAGCTGCCAGCGCAGCTTTCGCCAAAGCTTTTGTTCTAGTGCCAGT   774
sGFP2         ATCATGATATGGACGAGCTGCCAGCGCAGCTTTCGCCAAAGCTTTTGTTCTAGTGCCAGT   776
mNeonGreen    ATCATGATATGGACGAGCTGCCAGCGCAGCTTTCGCCAAAGCTTTTGTTCTAGTGCCAGT   780
vsfGFP-9      ATCATGATATGGACGAGCTGCCAGCGCAGCTTTCGCCAAAGCTTTTGTTCTAGTGCCAGT   753
NowGFP        ATCATGATATGGACGAGCTGCCAGCGCAGCTTTCGCCAAAGCTTTTGTTCTACTGCCAGT   777
muGFP         ATCATGATATGGACGAGCTGCCAGCGCAGCTTTCGCCAAAGCTTTTGTTCTAGTGCCAGT   776
mRFP1         ATCATGATATGGACGAGCTGCCAGCGCAGCTTTCGCCAAAGCTTTTGTTCTAGTGCCAGT   776
mOrange       ATCATGATATGGACGAGCTGCCAGCGCAGCTTTCGCCAAAGCTTTTGTTCTAGTGCCAGT   769
mStrawberry   ATCATGATATGGACGAGCTGCCAGCGCAGCTTTCGCCAAAGCTTTTGTTCTAGTGCCAGT   775
mApple        ATCATGATATGGACGAGCTGCCAGCGCAGCTTTCGCCAAAGCTTTTGTTCTAGTGCCAGT   776
mRuby2        ATCATGATATGGACGAGCTGCCAGCGCAGCTTTCGCCAAAGCTTTTGTTCTAGTGCCAGT   777
mScarlet-I    ATCATGATATGGACGAGCTGCCAGCGCAGCTTTCGCCAAAGCTTTTGTTCTAGTGCCAGT   768
mCherry       ATCATGATATGGACGAGCTGCCAGCGCAGCTTTCGCCAAAGCTTTTGTTCTAGTGCCAGT   769
FusionRedM    ATCATGATATGGACGAGCTGCCAGCGCAGCTTTCGCCAAAGCTTTTGTTCTAGTGCCAGT   777
```

Endogenous region Cytological location 68A4 11070449 ▼

```
mEmerald      GTTAGGCAGCATTTGAAATTTTTTGCCGGTGGATTGATTGTATTAAGGGGGGGGGGGGGG   838
mEGFP         GTTAGGCAGCATTTGAAATTTTTTGCCGGTGGATTGATTGTATTAAGGGGGGGGGGGGGG   838
sfGFP         GTTAGGCAGCATTTGAAATTTTTTGCCGGTGGATTGATTGTATTAAGGGGGGGGGGGGGG   834
sGFP2         GTTAGGCAGCATTTGAAATTTTTTGCCGGTGGATTGATTGTATTAAGGGGGGGGGGGGGG   836
mNeonGreen    GTTAGGCAGCATTTGAAATTTTTTGCCGGTGGATTGATTGTATTAAGGGGGGGGGGGGGG   840
vsfGFP-9      GTTAGGCAGCATTTGAAATTTTTTGCCGGTGGATTGATTGTATTAAGGGGGGGGGGGGGG   813
NowGFP        GTTAGGCAGCATTTGAAATTTTTTGCCGGTGGATTGATTGTATTAAGGGAGGGGGGGGGG   837
muGFP         GTTAGGCAGCATTTGAAATTTTTTGCCGGTGGATTGATTGTATTAAGGGGGGGGGGGGGG   836
mRFP1         GTTAGGCAGCATTTGAAATTTTTTGCCGGTGGATTGATTGTATTAAGGGGGGGGGGGGGG   836
mOrange       GTTAGGCAGCATTTGAAATTTTTTGCCGGTGGATTGATTGTATTAAGGGGGGGGGGGGGG   829
mStrawberry   GTTAGGCAGCATTTGAAATTTTTTGCCGGTGGATTGATTGTATTAAGGGGGGGGGGGGGG   835
mApple        GTTAGGCAGCATTTGAAATTTTTTGCCGGTGGATTGATTGTATTAAGGGGGGGGGGGGGG   836
mRuby2        GTTAGGCAGCATTTGAAATTTTTTGCCGGTGGATTGATTGTATTAAGGGGGGGGGGGGGG   837
mScarlet-I    GTTAGGCAGCATTTGAAATTTTTTGCCGGTGGATTGATTGTATTAAGGGGGGGGGGGGGG   828
mCherry       GTTAGGCAGCATTTGAAATTTTTTGCCGGTGGATTGATTGTATTAAGGGGGGGGGGGGGG   829
FusionRedM    GTTAGGCAGCATTTGAAATTTTTTGCCGGTGGATTGATTGTATTAAGGGGGGGGGGGGGG   837
```

Endogenous region Cytological location 68A4

```
mEmerald        GGGGGGG----------------------------------------   845
mEGFP           GGGGG------------------------------------------   843
sfGFP           GGGCNCCCCACAACGCAGGCATAATTAACGTGTCATACCTGTTCAAG   881
sGFP2           GGG--------------------------------------------   839
mNeonGreen      GGGG-------------------------------------------   844
vsfGFP-9        GGGGG------------------------------------------   818
NowGFP          G----------------------------------------------   838
muGFP           GG---------------------------------------------   838
mRFP1           GGGGGGGCCACCGGG--------------------------------   851
mOrange         GGGGGGGGGGGNNNNNN------------------------------   846
mStrawberry     GGGGGGGNNNGG-----------------------------------   847
mApple          G----------------------------------------------   837
mRuby2          GG---------------------------------------------   839
mScarlet-I      GGGGGGGGTG-------------------------------------   838
mCherry         GGGGGCGGCCGGGG---------------------------------   843
FusionRedM      GGG--------------------------------------------   840
```

**Figure S3:** Validation of landing sites for the H2Av:fluorophore insertion at attP2 sites in chromosome III. (A) PhiC31 recombination strategy used to generate transgenic H2Av:flurophore lines. Recombination between the attP site of chromosome III (attP2 site at 68A4 cytological location) of the landing line and the attB site of the plasmid containing the transgene in the presence of PhiC31 Integrase enzyme integrates the transgene into the specific attP (in this case att P2[2]) site. The attB and attP recombination generates attL and attR sites and the transgene is integrated in between. The attLFwd and 68A4 primers amplify 3' end of the transgene and the adjacent endogenous sequence. The sequence similarities of the endogenous sequence reveal the consensus in the background landing site of all the fluorophores. This strategy is used by Bestgene U.S.A. to confirm their insertions at attP2 site. (B) The multiple sequence alignment of the PCR amplified region from the genomic DNA between attL and 68A4 site. The amplified region includes the 3' end of the P-element (blue), and the endogenous region of the 68A4 location (orange). Note, the amplified endogenous region of 68A4 sequence (11070545—11070449) is similar across all the fluorophores insertion.

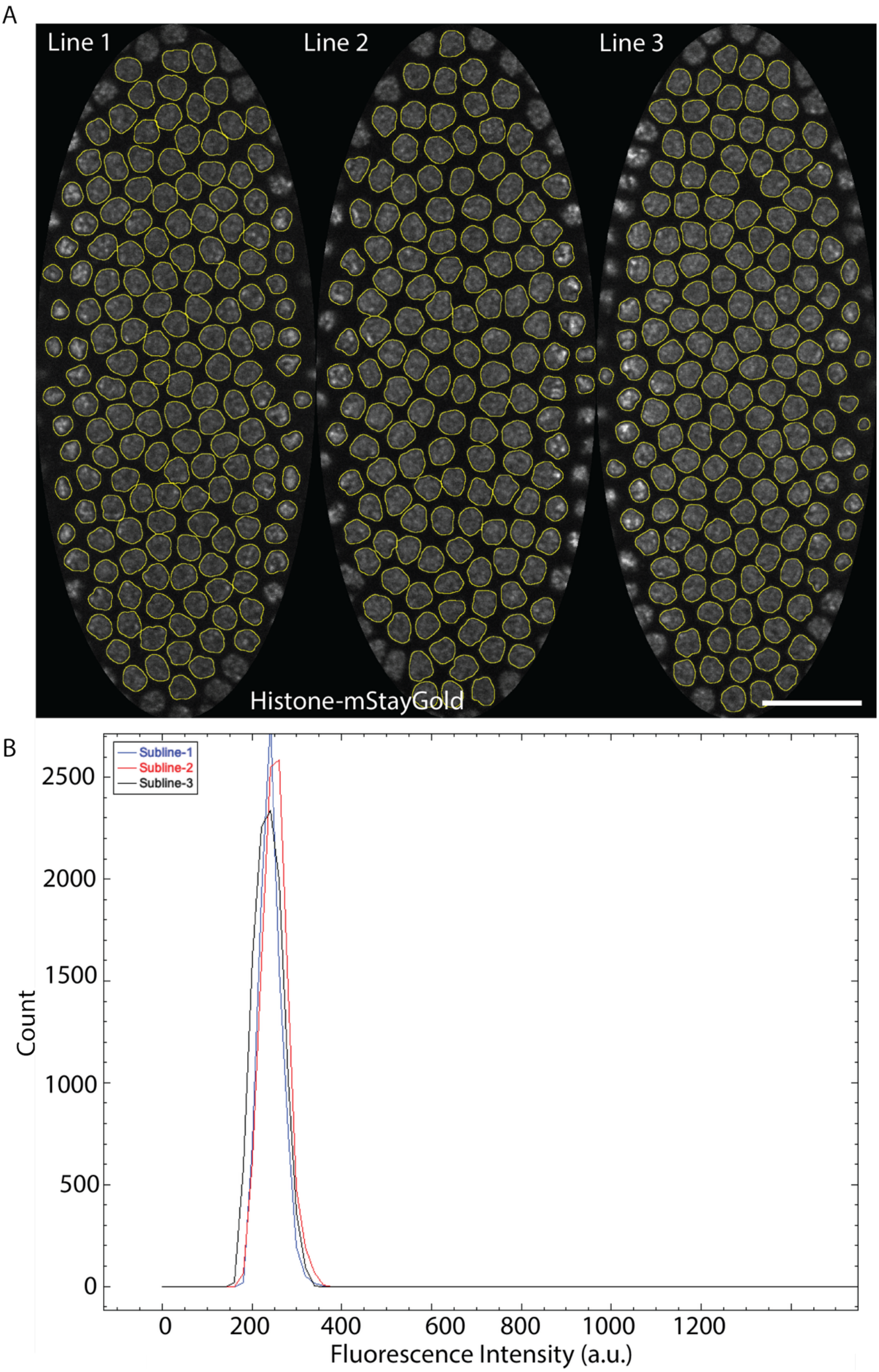


**Figure S4:** Comparison of three sublines for the H2Av::mStayGold insertion. (**A**) Comparison of the three sublines in n.c.14 of a single focal plane from representative embryos, with nuclei segmentation in yellow (scale bar = 20 μm). (**B**) Measured fluorescence intensity profiles. In this case, line 2 was selected for further analysis.

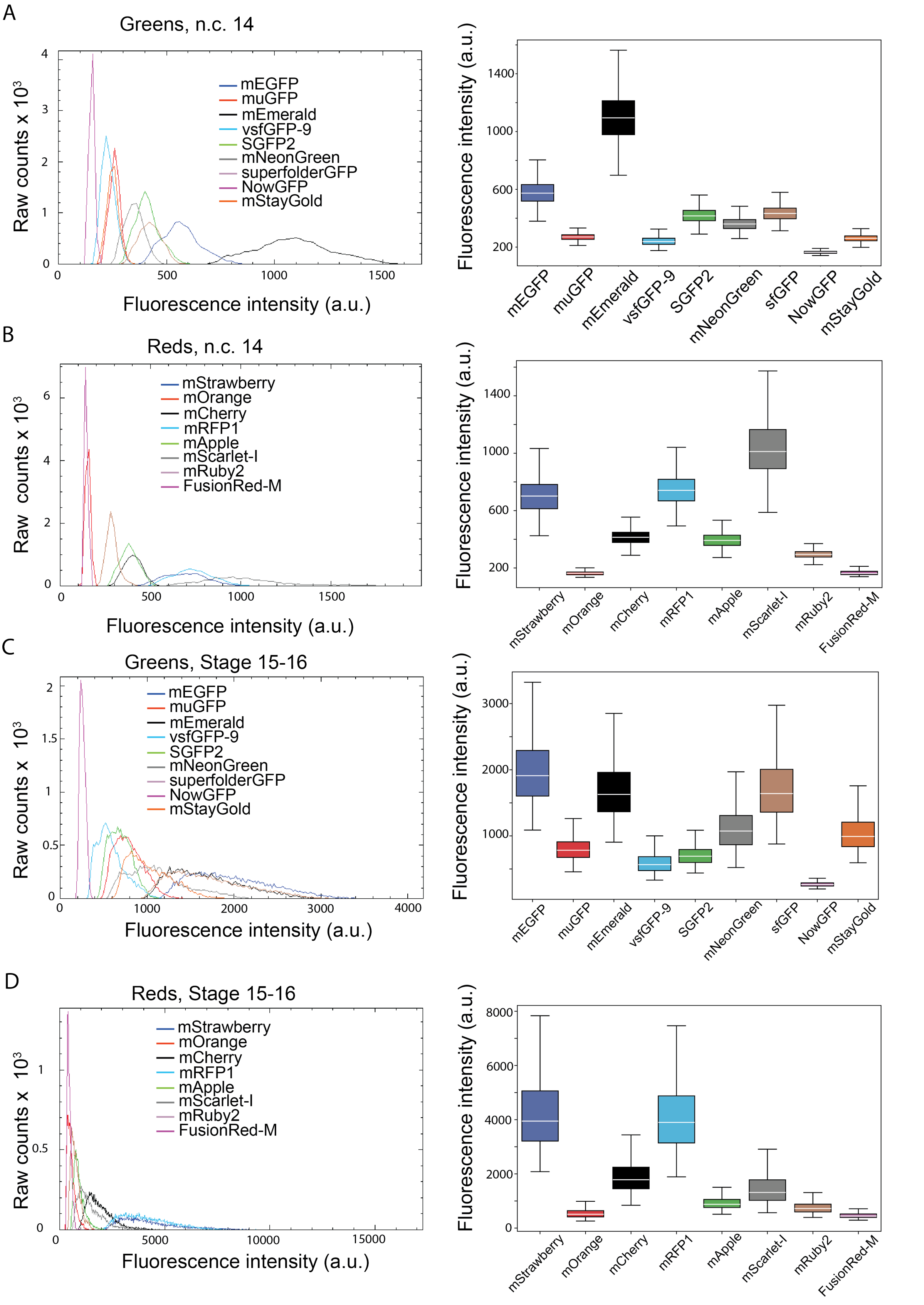


**Figure S5:** Raw measured intensity counts for the fluorophores in early (A-B) and late (C-D) embryos (left). Measured intensities for each fluorophore (right) showing median (horizontal bar) and boxes displaying the dataset quartiles, with the whiskers showing the spread in the distribution. At least three embryos per condition.

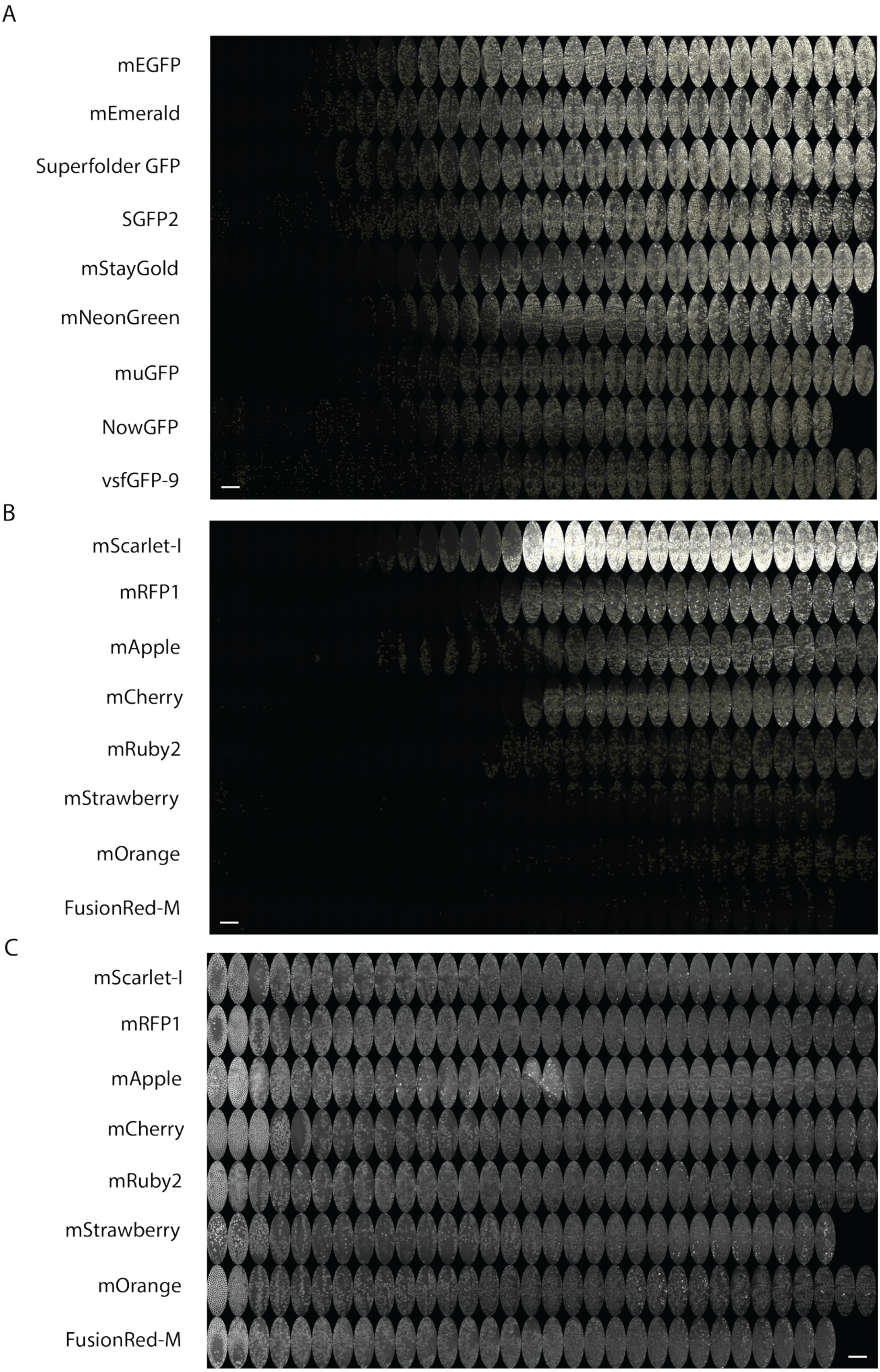


**Figure S6:** (**A-B**) Fluorophore folding time series for green (A) and red (B) fluorophores. Each image is 30 minutes apart and shows a single z-position. (**C**) Corresponding Nup::GFP signal for the images shown in (**B**). Scale bars = 50 μm.

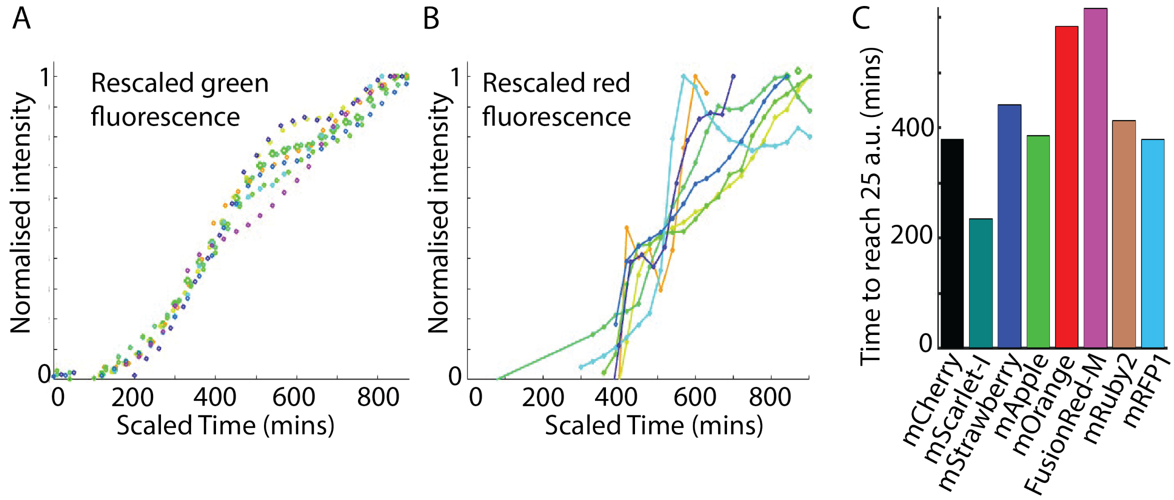


**Figure S7:** (**A-B**) Rescaled time for each fluorophore to align intensity profiles for green (A) and red (B) fluorophores. (**C**) Time to reach 25 a.u. for the red fluorophores. Data analysed is average of at least three embryos for each condition.

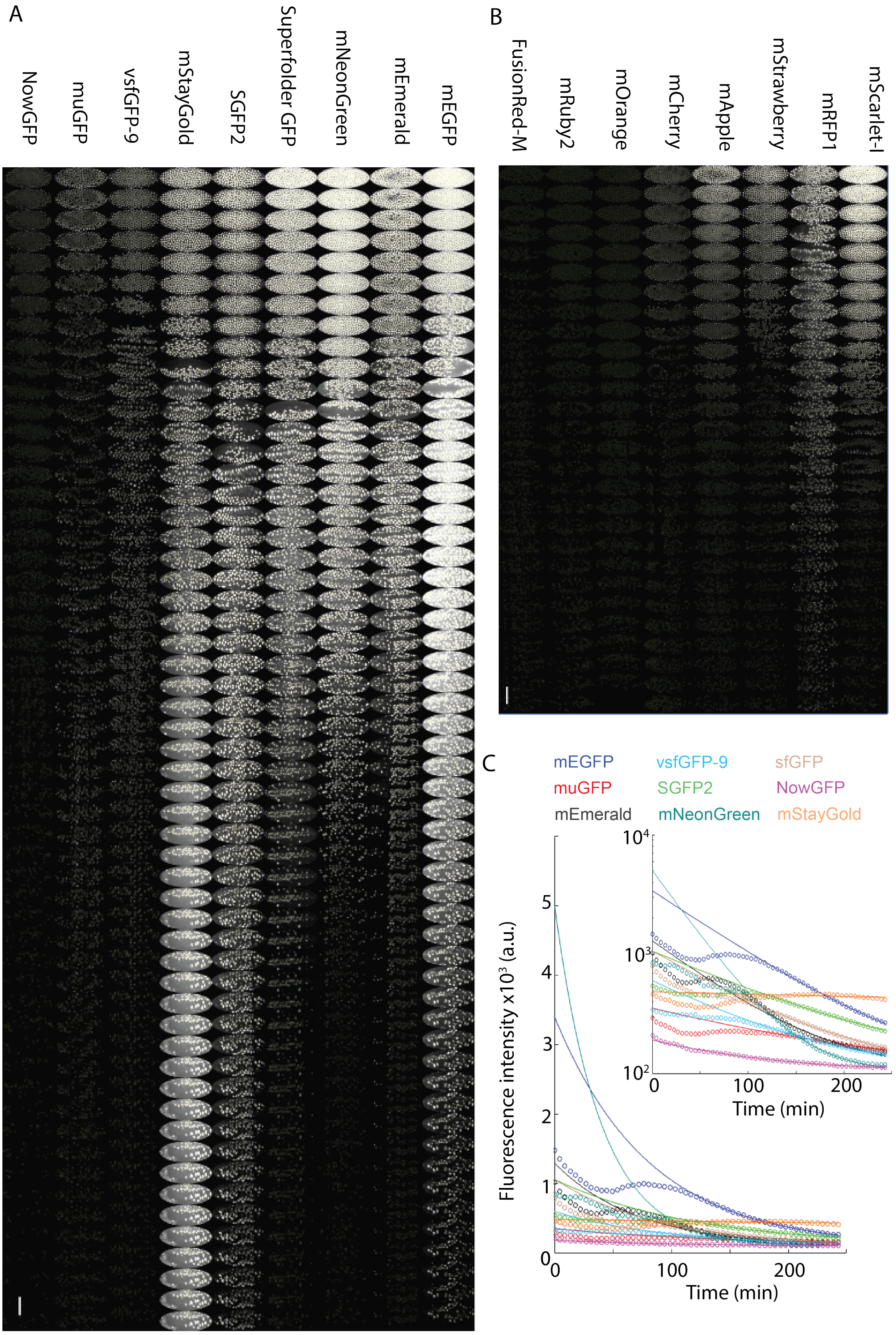


**Figure S8:** (**A-B**) Time course of bleaching for green (A) and red (B) fluorophores, showing a single z-stack position. Each time step is 4.5 mins. Scale bars = 50 μm. (**C**) Analysis of bleaching rate in green fluorophores, accounting for the intermediate increase in intensity. Inset same with logarithmic intensity scale. Fit to $I(t) = I_0 e^{-\alpha t} + I_1$ where $I_0, I_1$ and $\alpha$ are constants.

**Supplementary Movie Legends**

**Movie S1:** Z-stack through embryos expressing mEmerald (left) and mScarlet-I (right) in n.c. 14. The segmentation of nuclei is shown in yellow.

**Movie S2:** Folding - His2Av::mNeonGreen (green) and Nup::mRFP (magenta) embryo, starting in n.c. 14. Time step is 4.5 mins between frames.

**Movie S3:** Bleaching - His2Av::mEGFP embryo, starting in n.c. 14. Time step is 4.5 mins between frames.

**Movie S4:** Bleaching - His2Av::mStayGold embryo, starting in n.c. 14. Time step is 4.5 mins between frames.

**Supplementary Theory on Folding Rates**

By looking at the increase in signal over time defined from the beginning of n.c. 14 we can in principle estimate the fluorophore folding time. Histone mRNA is rapidly turned over in the *Drosophila* embryo, with a life time of approximately 20 mins around gastrulation[3]. The total histone mRNA rapidly reaches a plateau around 4-7 hours post oviposition[3]. We take the histone mRNA and protein (unfolded and folded) concentrations to be given by *m, U* and *M* respectively. We define *t = 0* as 30 minutes prior to gastrulation. mRNA is then produced at a rate *J* and degrades at rate $\sigma$. mRNA is then translated (assumed to be fast and hence neglected) at rate $\alpha$ and then folds at rate $\beta$ into a fluorescent protein. This protein degrades at rate $\mu$. This gives the following coupled equations:

$$\frac{dm}{dt} = J - \sigma m$$

$$\frac{dU}{dt} = \alpha m - \beta U - \mu U$$

$$\frac{dM}{dt} = \beta U - \mu M$$

These formulae are approximations. For example, the production of *mRNA J,* is likely to be both a function of space and time. In particular, this effect is clear in the green fluorescence data (the intermediate peak in intensity seen in Fig. S8C).

These equations can be solved exactly, giving the histone protein with unfolded fluorophore as:

$$m(t) = \frac{J}{\mu}(1 - e^{-\sigma t})$$

$$U(t) = \frac{J}{\mu}\left(\frac{1 - e^{-(\beta+\mu)t}}{\beta + \mu} - \frac{e^{-\sigma t} - e^{-(\beta+\mu)t}}{\beta + \mu - \sigma}\right)$$

and similar for M(t).

However, this form is not so useful as relies on many assumptions. Therefore, we assume that the mRNA equilibrates much faster than the protein, so

$$\frac{dU}{dt} = g - (\beta + \mu)U$$

Therefore, $U = \frac{g}{\beta+\mu}\left(1 - e^{-(\beta+\mu)t}\right)$. Now

$$\frac{dM}{dt} = \frac{g\beta}{\beta + \mu}\left(1 - e^{-(\beta+\mu)t}\right) - \mu M$$

Assuming a linear relationship between concentration and observed fluorescence ($\phi_f$), then the observed intensity *I(t)* follows an exponential like curve. As seen from Fig. 3-4, the observed fluorescence varies between fluorophores. This idealised approach was not particularly helpful in the *in vivo* context, due to the large variability between fluorophores and that protein production was clearly not constant in time. Therefore, we utilised more phenomenological approaches to assay the folding rates as described in the Methods.
